# Systematic analysis of disease-linked rare germline variants reveals new classes of cancer predisposing genes

**DOI:** 10.1101/2023.02.15.528610

**Authors:** Seulki Song, Youngil Koh, Seokhyeon Kim, Sang Mi Lee, Hyun Uk Kim, Jung Min Ko, Se-Hoon Lee, Sung-Soo Yoon, Solip Park

## Abstract

Despite recent insights into cancer predisposition genes (CPGs), the contribution of rare germline variants to carcinogenesis remains unclear. Here, we examine the enrichment of rare germline variants in Mendelian disease-associated genes using PCAWG and the 1,000 Genomes Project. Using an integrative approach that considers disease classes, cellular pathways, and tissue expression profiles, we find that subsets of Online Mendelian Inheritance in Man (OMIM) genes are strong candidate CPGs. These genes were classified into five clusters suggesting diverse mechanisms underlie tumor progression. We further analyzed *PAH* gene which shows the strongest prevalence of rare germline variants in cancers (2.2%) compared to controls (0.7%). This enrichment was validated in two independent cancer cohorts and present the possibility of tumorigenesis by contributing immune-associated pathways. Our data argue that rare germline variants of Mendelian disease-associated genes contribute to cancer progression and suggest new CPG classifications that may underlie diverse tumorigenesis mechanisms.

## Introduction

Germline variants can lead to an increased risk of cancer in individuals. The existence of such a genetic predisposition to cancer was first proposed by Broca who noted breast cancer in 15 members of his wife’s family ^1^. Since this seminal finding, approximately 140 CPGs have been discovered through multiple strategies that include genome-wide linkage analysis with small family studies to large-scale next generation sequencing ^2^. However, there may be many genes associated with cancer predisposition that have not been identified. This is due to the low statistical power to detect associations between rare pathogenic germline variants and cancer risk ^2^.

In addition to difficulties with identifying CPGs, an understanding of their functional contribution to cancer is still limited. However, they are known to be closely linked to fundamental cellular processes which are core to the function of cancer cells ^3^. Furthermore, CPGs are typically monogenic and show high-penetrance phenotypes and bi-allelic inactivation in tumors ^2, 4^. Currently, knowledge from large-scale cancer consortium data indicates possible novel mechanisms of CPGs that include immune evasion, epigenetic reprogramming, and sustained proliferative signaling also contribute to carcinogenesis ^5^. Consistent with this, large-scale tumor sequencing data has led to the systematic evaluation of whether diverse genes beyond CPGs increase the risk of cancer ^6^.

To systemically identify novel CPGs, we focused on Mendelian disease-associated genes from Online Mendelian Inheritance in Man (OMIM) which is a catalog of human genes associated with heritable disorders that are mainly identified by genetic linkage studies^7^. We reasoned that disease-associated genes in OMIM are similar to CPGs in that they are monogenic and show high-penetrance phenotypes. Indeed, it has been reported that patients with genetic disorders linked to rare variants in OMIM genes show secondary phenotypes in adulthood. For example, patients with rare variants in *GBA* genes, which are associated with Gaucher’s disease in childhood, present with Parkinson’s disease as adults ^8^. These examples indicate that variants in OMIM genes that are known to be associated with Mendelian diseases may also contribute to additional disorders including cancer.

In this study, we evaluated OMIM genes as potentially novel CPGs. We tested (i) whether there is an enrichment for pathogenic germline variants of OMIM genes in cancer patients compared to control; and (ii) if so, whether OMIM genes could be classified by integrating multiple features to represent diverse CPG classes. After case-control analysis, two features were also considered to define diverse classes of OMIM genes as potential CPGs, prevalence of bi-allelic inactivation and gene expression profile across tissues.

Canonical CPGs such as *BRCA1/2* are ubiquitously expressed non-specifically in many tissues ^2^. Furthermore, they tend to follow the ‘two-hit’ hypothesis where a pathogenic germline variant occurs in one allele of the gene and a ‘second hit’ occurs somatically in the second allele leading to increased cancer risk ^2^. However, exceptions to broad tissue expression are discovered in several CPGs ^2^ and furthermore, diverse mechanisms beyond the canonical CPG model are being suggested ^6, 9^. For example, pathogenic variants in α-1-antitrypsin gene, *SERPINA1,* increase susceptibility to breast cancer ^10^, although *SERPINA1* is preferentially expressed in the liver (at the RNA level) or in the lung, kidney, and gastrointestinal tract (at the protein level) ^11^. Furthermore, some CPGs show that only single inherited variants can suffice to increase cancer risk. We hypothesized that heterozygote pathogenic germline carriers might also develop cancer through mechanisms beyond the classical two-hit hypothesis.

In this study, we assumed that rare pathogenic germline variation in OMIM genes, such as genetic haploinsufficiency in hereditary diseases, will also increase the risk of tumor development. To verify this hypothesis, we performed a comprehensive analysis using the large-scale cancer genomics data from PCAWG and the 1000 Genome project as healthy control. Our results suggest that multiple OMIM genes also harbor pathogenic germline variants that predispose individuals to cancer in the general population. Furthermore, we deeply analyzed the tumorigenesis mechanism of *PAH*, which presents the highest pathogenic germline variant frequencies in OMIM genes, and uncovered novel mechanisms where OMIM genes increase cancer risk by modulating metabolic and immune response-associated pathways.

## Result

### A framework for measuring enrichment of germline variants in cancer

To systematically estimate the contribution of rare pathogenic germline variants in OMIM genes to cancer, we designed a statistical method that tests the excess of rare pathogenic germline variants in cancer patients compared to healthy control samples.

Our approach was intended to collect only high-confidence pathogenic variants. The potential pathogenic variants were first selected if they have a minor allele frequency (MAF) < 0.5% in the Genome Aggregation Database (gnomAD) exome ^12^ (see **Methods**). Next, we considered three sets of possible pathogenic variants with stringent variant curation steps (i) Tier1: protein truncation variants (PTVs, encompassing splicing variants, frameshift indels, and nonsense variants) and (ii) Tier2: previously reported to be clinically significant (designated “pathogenic” or “likely pathogenic”) in the ClinVar database ^13^ using the clinical standard ACMG/AMP criteria (see **Methods**). (iii) Overall Tier1+2: either Tier1 or Tier2 incorporating to estimate the maximal contribution of possible pathogenic variants to cancers.

We aggregated single nucleotide variants (SNVs) and indels in each gene in order to examine the contribution of selected pathogenic variants to cancer risk. The case-control method tests for an excess of pathogenic variants in a gene in the case (cancer patients) compared to the control (non-cancer patients). This analysis uses a burden test to control for population structures with the first two PC values based on common variants of case-control samples (**Fig. 1a, Supplementary Fig. 1a-b**; see **Methods**).

**Figure 1.**
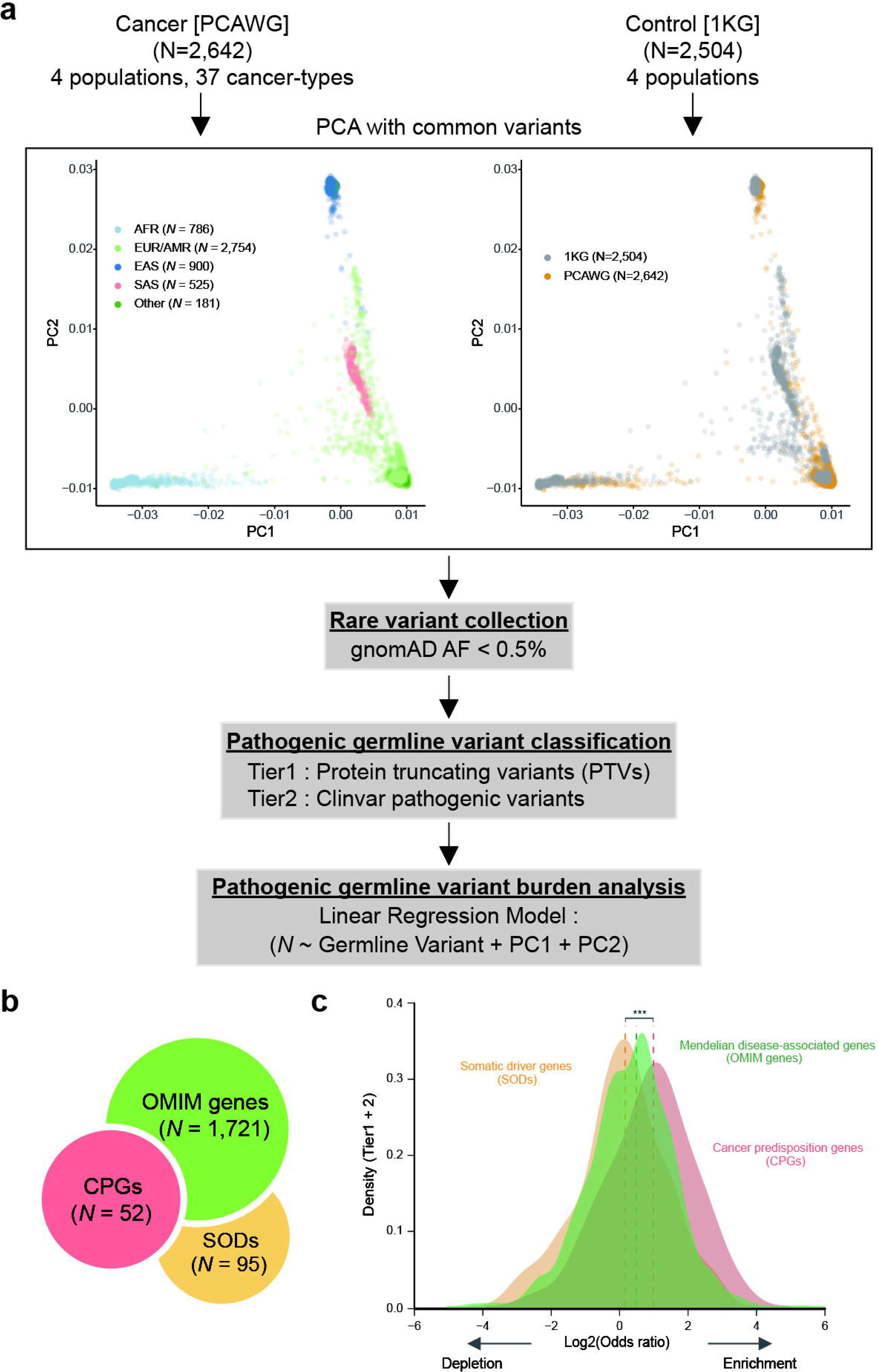
Systematic analysis for the enrichment of rare pathogenic germline variants in cancer cases compared to control samples. **(a)** Overview of case-control analysis. Principal components analysis (PCA) using common variants was performed to adjust the population of cancer patients and control individuals. After collecting rare pathogenic germline variants from case-control cohorts, the linear regression model tests germline variant enrichments in cancer cases compared to control with the first two PC values used to stratify the population. AFR: African, AMR: American, EAS: East Asian, EUR: European, and SAS: South Asian. **(b)** Genes with at least 3 carriers were tested in this study including cancer predisposition genes (CPGs), somatic driver genes (SODs), and Online Mendelian Inheritance in Man genes (OMIM genes). **(c)** Enrichment of rare pathogenic germline variants (Overall: Tier1 and 2) in cancer compared to control across three gene sets (* *P* < 0.05, *** *P* < 0.001).

### Rare germline variant enrichment of Mendelian disease-associated genes in Pan-cancer

We collected two distinct data sets for which individual-level genome sequences were available. The first dataset is a compilation of multi-ancestral studies, involving 2,642 cancer patients across four subpopulations from the Pan-Cancer Analysis of Whole Genomes Network (henceforth PCAWG) ^14^. The second dataset includes 2,504 healthy control exomes across four subpopulations from the 1000 Genomes Project (1KG) ^15^ (see **Methods**; **Supplementary Fig. 1c, Supplementary Table 1**). The frequency of pathogenic variants (either Tier1 or Tier2) varied widely across samples and also cancer types with 25,099 pathogenic variants (16,259 for case patients and 11,332 for control samples) from 8,397 genes that have at least one pathogenic variant. To increase statistical power, we restricted the regression analysis to genes with at least 3 carriers (*N* = 4,705 genes; **Fig. 1b-c** and **Supplementary Table 2**).

In a Pan-cancer analysis, we first observed that previously known cancer predisposition genes (CPGs; *N*=52) based on the recent literature review ^16^ (**Fig. 1b**) showed a significant enrichment of Tier1 variants in cases compared to controls (median Log2 odds ratio (OR) for the enrichment of Tier1 in cases compared to controls = 1.00, *P* = 3.73 × 10^−4^ by Wilcoxon rank sum test, **Fig. 2a**). Above all, 6 CPGs (11.5 %) were significantly enriched for Tier1 in cancer cases versus controls individually (*P* < 0.05 by Pan-cancer case-control analysis; **Supplementary Table 2**). For example, *BRCA2* showed the highest cancer prevalence compared to controls (Log2 OR = 3.46, *P* = 1.64 ×10^−3^) and other significant CPGs including *XPA* (Log2 OR = 3.18, *P* = 4.64 × 10^−2^), *POLH* (Log2 OR = 2.68, *P* = 9.82 × 10^−7^), *BRCA1* (Log2 OR = 2.62, *P* = 2.51 × 10^−3^), *FAH* (Log2 OR = 2.52, *P* = 4.17 × 10^−2^), and *ATM* (Log2 OR = 1.67, *P* = 3.38 × 10^−2^).

**Figure 2.**
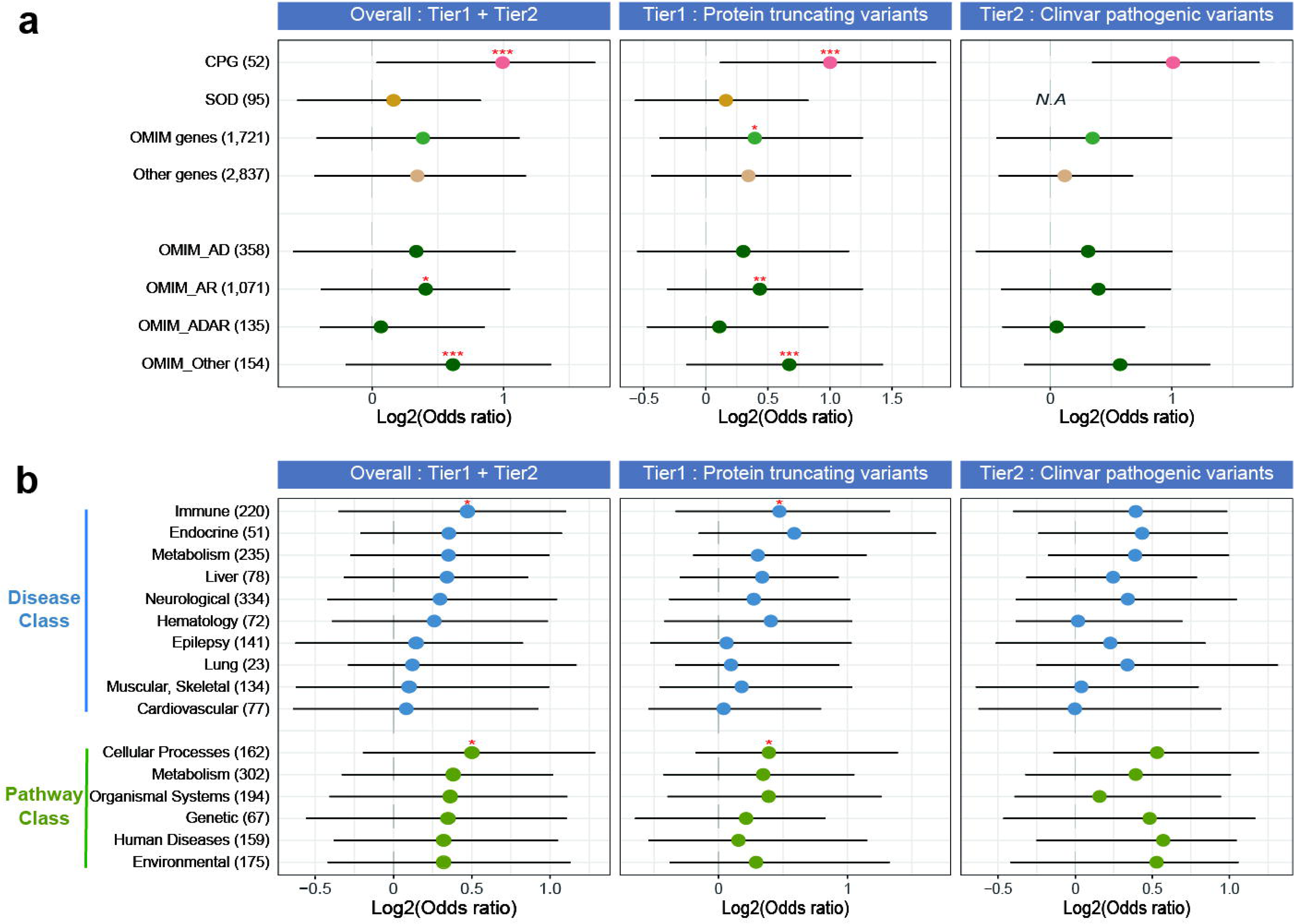
Enrichment of rare pathogenic germline variants in 2,642 cases over 2,504 control samples. (a) Log2 odds ratio distribution with case-control analysis across four different gene sets and four inheritance groups of OMIM genes (* *P* < 0.05, ** *P* < 0.01, *** *P* < 0.001). The length of each whisker is 1.5 times the interquartile range (shown as the height of each box). **(b)** Log2 odds ratio distribution from case-control analysis across disease classes and cancer-related pathways using OMIM genes. Dots for each group represent the median value of the prevalence for each group (* *P* < 0.05). The length of each whisker is 1.5 times the interquartile range (shown as the height of each box).

We next tested whether Mendelian disease-associated genes in OMIM (OMIM genes; *N*=1,721) that contribute to genetic diseases but have not previously been associated with cancer risk show enrichment of germline variants in cancer patients compared to control samples (**Fig. 1b**). Notably, OMIM genes also showed significant enrichment for Tier 1 variants in cases compared to controls (median Log2OR = 0.39, *P* = 1.64 × 10^−2^ by Wilcoxon rank sum test, **Fig. 2a**). This enrichment was stronger when applying OMIM sub- groups separately, OMIM-Autosomal Recessive genes (OMIM-AR; median Log2OR = 0.43, *P* = 8.34 × 10^−3^ by Wilcoxon rank sum test, **Fig. 2a**) and OMIM genes with X or Y-linked, somatic or multifactorial inheritance (OMIM-Other; median Log2OR = 0.66, *P* = 2.96 × 10^−3^ by Wilcoxon rank sum test, **Fig. 2a**). 59 OMIM genes (3.4%) were individually significantly enriched for Tier1 in cases versus controls (*P* < 0.05 by Pan-cancer; **Supplementary Table 2**) including *FYCO1* (Log2 OR = 4.08, *P* = 1.61 × 10^−2^), *PLOD3* (Log2 OR = 3.70, *P* = 2.04 × 10^−2^), *CD96* (Log2 OR = 3.67, *P* = 1.72 × 10^−2^), *NR1H4* (Log2 OR = 3.66, *P* = 3.84 × 10^−2^), and *ALB* (Log2 OR = 3.51, *P* = 2.54 × 10^−2^). This indicates that rare pathogenic germline variants in many OMIM genes may also contribute to increased cancer risk.

As a control gene set, we considered somatic driver genes (SODs; *N*=95) that are known to increase cancer risk by frequent somatic alterations based on the integration of three different sources (see **Methods, Fig. 1b**). We found no cancer enrichment in these genes (median Log2OR = 0.16, **Fig. 2a**). A consistent result was obtained when we applied non-cancer related genes (Others; *N*=2,837; median Log2OR = 0.34, *P* = 6.17 × 10^−2^ by Wilcoxon rank sum test**, Fig. 2a**). It should be noted that this yields a conservative upper limit since the non-cancer related genes may also include possible cancer predisposition genes. Our analysis can, therefore, specifically provide the evidence of OMIM genes as possible CPGs, which show significant enrichments of germline variants in cancer compared to control samples.

In the Tier2 variants, CPGs showed the strongest enrichment in a Pan-cancer (median Log2 OR = 1.01). OMIM genes also showed continually high enrichment (median Log2 OR = 0.35) but SODs did not have any clinically proven pathogenic variants from ClinVar ^13^. Overall, Tier1 and 2 variants presented similar enrichment results in a Pan-cancer. CPGs presented the strongest enrichment (median Log2 OR = 0.99; *P* = 2.64 × 10^−4^ by Wilcoxon rank sum test) next to OMIM genes (median Log2 OR = 0.39; *P* = 5.45 × 10^−2^ by Wilcoxon rank sum test) including OMIM-AR (median Log2 OR = 0.41; *P* = 3.85 × 10^−2^ by Wilcoxon rank sum test) and OMIM-Other (median Log2 OR = 0.61; *P* = 4.41 × 10^−3^ by Wilcoxon rank sum test). Since this provided the maximal contribution of presumptive pathogenic germline variants, we further analysed using Tier1 and 2 variants as pathogenic germline variants (**Fig. 2a**).

### Rare germline variant enrichment of OMIM genes across single-cancers

We also applied case-control analysis to each of the 26 cancer types with > 20 samples (95.3% of the samples in a Pan-cancer; **Supplementary Fig. 2, Supplementary Table 3**). At the single gene level, 165 genes (9 CPGs and 156 OMIM genes with 242 gene-cancer associations) were significantly associated with at least one individual cancer type and 50 genes were also significantly enriched in a Pan-cancer (*P* < 0.05; **Supplementary Table 2**). In CPGs, as expected, *BRCA1* was significantly enriched in ovarian adenocarcinoma (Ovary-AdenoCA; Log2OR = 8.57, *P* = 6.28 × 10^−1^^2^) and breast adenocarcinoma (Breast-AdenoCA; Log2OR = 4.35, *P =* 6.39 × 10^−4^; **Supplementary Table 2**), and *BRCA2* in Ovary-AdenoCA (Log2OR = 7.16, *P* = 2.35 × 10^−6^), Pancreatic Adenocarcinoma (Panc-AdenoCA; Log2OR = 5.98, *P* = 1.81 × 10^−6^) and Breast-AdenoCA (Log2OR = 5.55, *P* = 3.84 × 10^−5^; **Supplementary Table 2**). Among OMIM genes, we identified: *GABRA1* (Log2OR = 57.73, *P* = 2.33 × 10^−2^^2^) in chronic lymphocytic leukemia (Lymph-CLL) was the top ranked significantly enriched gene-cancer pair. In addition, *PEX1* (Log2OR = 43.73, *P* = 3.17 × 10^−1^^6^) in B-cell non-Hodgkin’s lymphoma (Lymph-BNHL), *FAM111A* (Log2OR = 21.19, *P* = 1.27 × 10^−1^^2^) and *CD96* (Log2OR = 21.19, *P* = 1.39 × 10^−1^^2^) in medulloblastoma (CNS-Medullo), and *SLC24A4* (Log2OR = 16.92, *P* = 1.29 × 10^−10^) in liver hepatocellular carcinoma (Liver-HCC) also showed significant enrichment.

### Germline variant enrichment based on disease class

Many OMIM genes that cause different genetic diseases can also be grouped into disease classes based on the physiological system affected. To complement single-gene-based case-control analysis, we measured the enrichment of pathogenic germline variants (overall Tier1 +2) in each disease class instead of single diseases. From 93 genetic diseases with 1,721 OMIM genes, 10 representative disease classes were defined based on the specific physiological system affected by disease (see **Methods**). The immune disease class (including 220 OMIM genes from 12 diseases) showed the strongest enrichment in a Pan-cancer (median Log2OR = 0.47, **Fig. 2b**). This enrichment is driven by rare pathogenic variants including *G6PD* (Log2OR = 5.30, *P* = 5.64 × 10^−4^), *C7* (Log2OR = 1.55, *P* = 6.41 × 10^−3^), *MEFV* (Log2OR = 1.87, *P* = 9.79 × 10^−3^) and *PHYH* (Log2OR = 2.20, *P* = 1.16 × 10^−2^). Next, the endocrine disease class (including 51 OMIM genes from 7 diseases) and metabolic class (including 235 OMIM genes from 27 diseases) also showed strong enrichment of pathogenic variants (median Log2OR =0.35 for both classes). This included *PAX4* (Log2OR = 2.86 *P* = 1.22 × 10^−4^) in the endocrine disease class and *CYP24A1* (Log2OR = 2.13, *P* = 7.38 × 10^−3^), SLC22A12 (Log2OR = 1.68, *P* = 9.28 × 10^−3^), *PHYH* (Log2OR = 2.20, *P* = 1.16 × 10^−2^), *NPC2* (Log2OR = 1.67, *P* = 1.21 × 10^−2^), and *ACADM* (Log2OR = 1.57, *P* = 1.24 × 10^−2^) in the metabolic disease class.

We next performed the disease class-based enrichment analysis on each of the 26 cancer types (**Fig. 3a** and **Supplementary Table 4**). Overall, most tested genetic diseases for 10 representative disease classes (63.4%; 59 out of 93 at *P* < 0.05) showed significant enrichment in at least one single cancer type or Pan-cancer. The immune disease class had the highest number of significantly enriched cancer types (80.8%; 21 out of 26 cancer types at *P* < 0.05). The neurological disease class had the second- highest numbers (76.9%; 20 cancer types) followed by the metabolic disease class (57.7%; 15 cancer types). In detail, common variable immune deficiency (CVID) from the immune disease class in bladder transitional cell carcinoma (Bladder-TCC) showed the strongest enrichment (Log2OR = 8.26, *P* = 1.43 × 10^−2^). CVID is known to be associated with an increased risk of malignancy and tumorigenesis ^17, 18^. Moreover, holoprosencephaly in the neurological disease class of biliary adenocarcinoma (Biliary-AdenoCA; Log2OR = 7.66, *P* = 1.27 × 10^−2^) and lipodystrophy in the metabolic class of Lymph-BNHL (Log2OR = 6.45, *P* = 6.02 × 10^−4^) showed next significant enrichments. Additionally, high bone density disorders from the immune disease class were strongly enriched in lung squamous cell carcinoma (Lung-SCC; Log2OR = 5.39, *P* = 3.45 × 10^−2^). Interestingly, the comorbidity relationship between bone mineral density disorder and lung cancer has previously been observed ^19, 20^.

**Figure 3.**
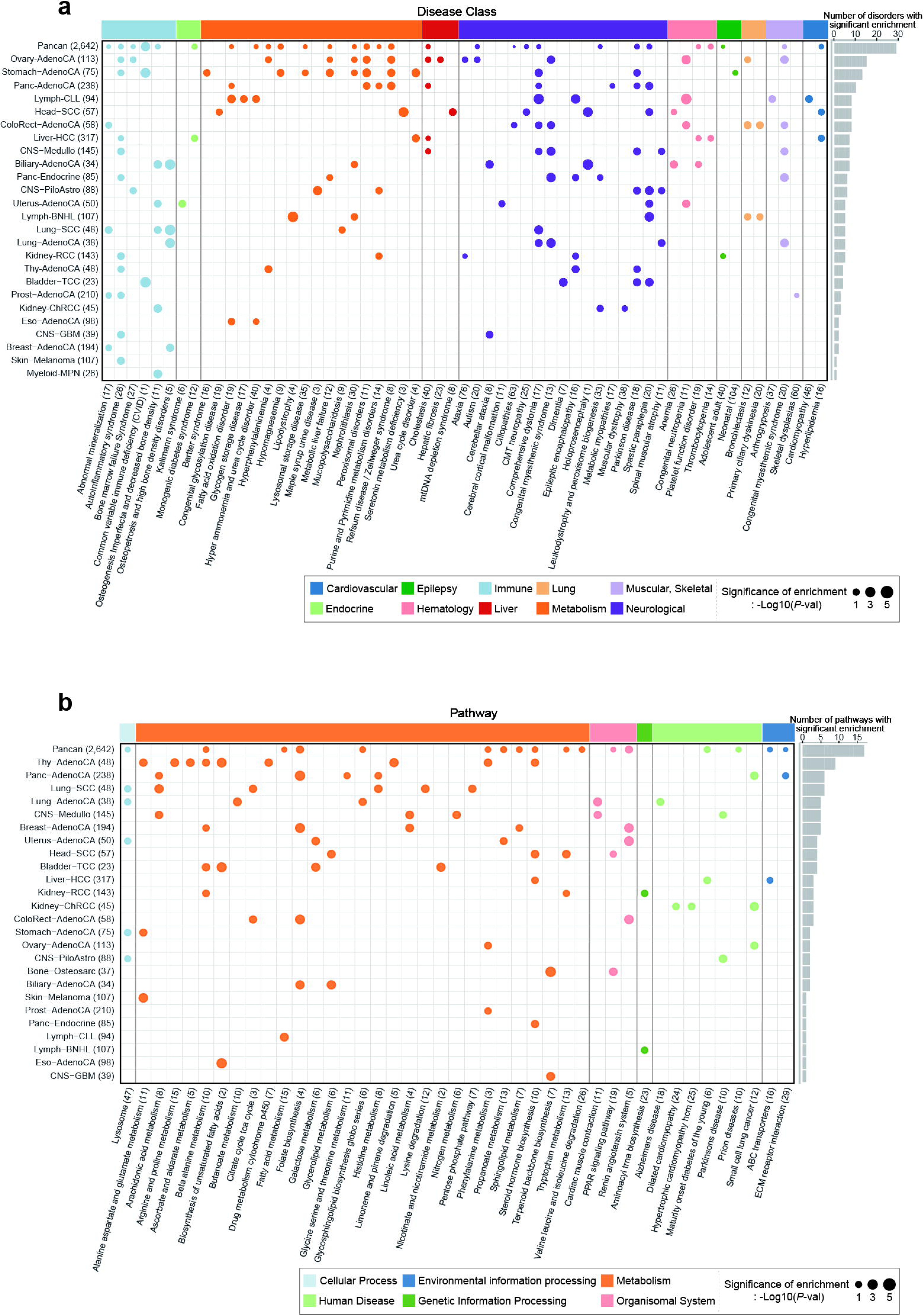
Pathogenic germline variants enriched in (a) disease classes and (b) KEGG pathways across Pan-cancer and single cancer types using linear regression model after population adjustment. Circle size indicates the significance of pathogenic germline variant enrichment in a cancer type compared to control and colour represents a disease class (or KEGG pathway). Only single cancer types with > 20 samples are presented. Numbers in parentheses indicate number of tested samples in each cancer type (left) and tested OMIM genes in each disease (or KEGG pathway; bottom). The number of detected diseases (or KEGG pathways) (at *P* < 0.05) in each cancer type are presented in the bar plot.

We also performed deeper enrichment analysis using 70 KEGG signalling pathways to examine the cancer-associated OMIM genes identified by our analysis compared to controls (see **Methods**; **Fig. 3b** and **Supplementary Table 4**) ^21^. 43 pathways showed significant enrichment of pathogenic germline variants in at least one individual cancer type and Pan-cancer compared to controls (61.4%, 43 out of 70; *P*-value < 0.05). The metabolic pathway class (including 29 pathways) showed the highest number of significantly enriched cancer types (88.5%; 23 out of 26 cancer types) followed by the organismal system pathway class (34.6%; 9 out of 26 cancer types with 3 pathways) and human disease pathway class (30.8%; 8 out of 26 cancer types with 7 pathways). Specifically, genes associated with the terpenoid backbone biosynthesis pathway showed the strongest enrichment in osteosarcoma (Bone-Osteosarc) (Log2OR = 5.32, *P* = 1.91 × 10^−2^). The second strongest enrichment, unsaturated fatty acid biosynthesis in esophagus adenocarcinoma (Eso-AdenoCA, Log2OR = 5.16, *P* = 1.48 × 10^−2^).

### Five possible classes in OMIM genes

To systematically investigate how OMIM genes contribute to increased cancer risk, we integrated four features: (i) prevalence of germline variants in cancer compared to control (case-control analysis), (ii) two-hit preference, (iii) overall gene expression levels across tissues and (iv) tissue-specificity score. We expected to observe diverse classification of OMIM genes based on the relatedness of CPGs by applying these four features.

We applied the Knudson’s two-hit hypothesis across each gene in a Pan-cancer. The two-hit preference in this study was designed to measure the enrichment of pathogenic germline variants (overall Tier1 +2) in samples with loss of heterozygosity (LOH) compared to samples without LOH within a gene (see **Methods**). Next, we applied two gene expression level-based features from the Human Protein Atlas (HPA) ^22^: overall properties of gene expression level across 20 tissues (the number of tissues which are expressed higher than the mean expression value of each gene) and tissue specificity (number of highly expressed tissues above 2 normalised expression level). Although many CPGs are known to be ubiquitously expressed, there is diversity in tissue-specific expression for some CPGs (**Supplementary Fig. 3**).

To improve statistical power, our analysis was restricted to genes with consistent enrichment of pathogenic germline variants in cancer patients compared to control samples not only in PCAWG, but also in independent cancer cohort (TCGA after removing the overlapping samples with PCAWG, see **Methods; Fig. 4a-b,** and **Supplementary Fig. 4a**). Next, we selected genes with pathogenic variants that met our threshold (at least 3 carriers in overall Tier1 and 2 in PCAWG). Ultimately, 321 genes (including 17 CPGs and 304 OMIM genes) were selected (**Fig. 4b**).

**Figure 4.**
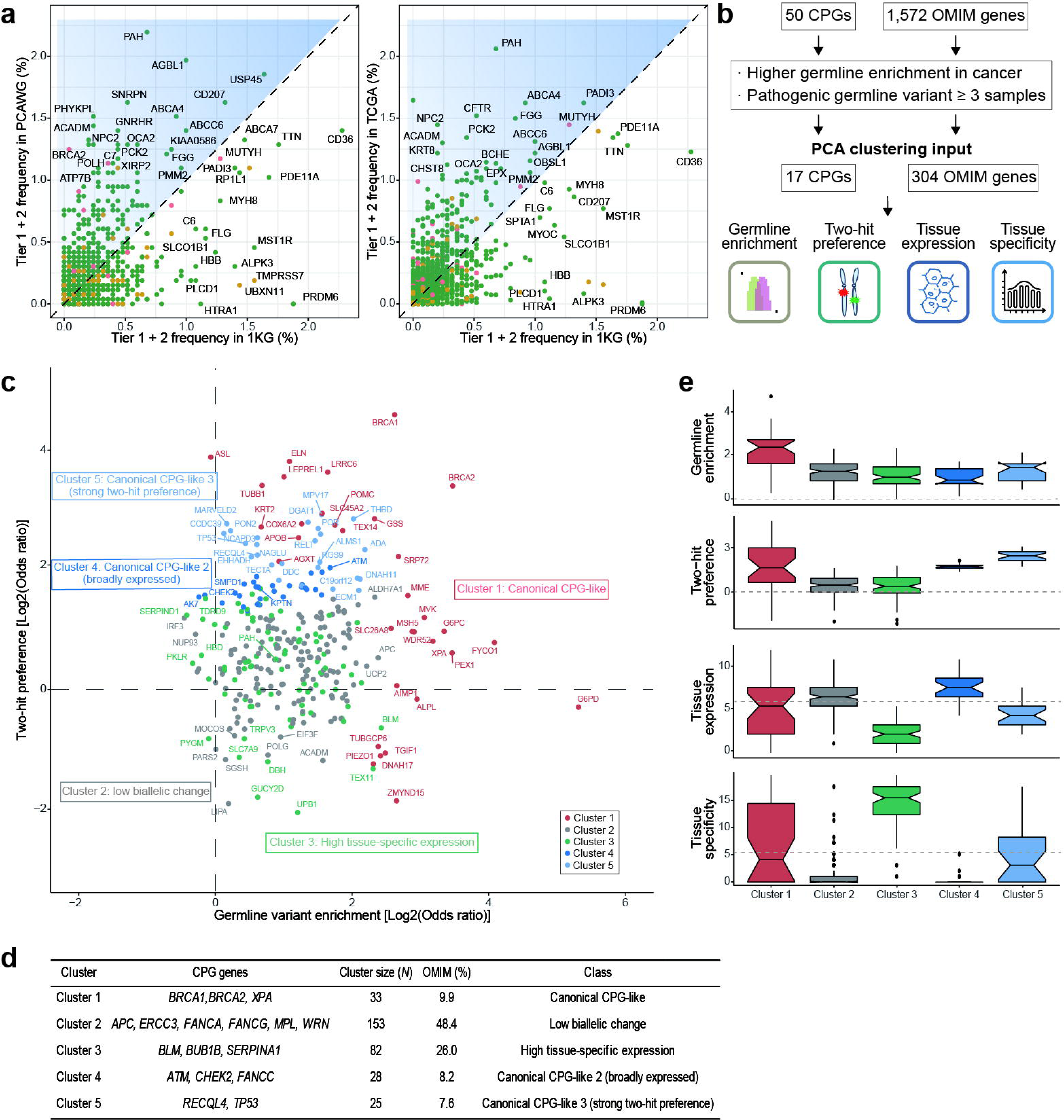
Gene classifications for CPGs and OMIM genes. (a) Genes with high prevalence of pathogenic germline variants in cancer (both in PCAWG and TCGA) compared to control were selected. The color indicates type of gene set (red: CPGs, green: OMIM genes, and yellow: SODs, respectively). **(b)** Schematic representation of gene classification by integrating four features: (i) Germline enrichment in cancer: pathogenic germline variant enrichment in case compared to control, (ii) two-hit preference: excess of pathogenic germline variants in samples with LOH compared to samples without LOH, (iii) tissue expression: number of tissues have higher expression level of mean value across 20 different tissues, and (iv) tissue-specificity: number of highly expressed tissues above 2 normalized expression. **(c)** Scatter plot presenting rare germline variant enrichments (x-axis) and two-hit preferences (y-axis) of CPGs and OMIM genes in a Pan-cancer. Each gene is coloured by assigned gene cluster. **(d)** Summary table for each class. **(e)** Box plot presenting the characteristics of each cluster. Tissue specificity indicates the total tissue types (*N* = 20) minus the number of tissues with tissue expression greater than 2 normalized expressions. More positive values in tissue expression indicate broader expression across tissue types and more positive values in tissue specificity indicate higher tissue-specific gene expression. The circles indicate outliers, which are values between 1.5 and 3 times the inter-quartile ranges. The first two horizontal dashed lines of germline enrichment and two-hit enrichment indicate a Log2(odds ratio) equal to zero. The next two grey dashed lines indicate the median value of tissue expression and tissue specificity.

We performed principal component analysis (PCA) by applying the above four features to input genes. Overall, we defined five clusters after optimization using a trimmed k-means clustering algorithm, suggesting that diverse mechanisms predispose OMIM genes to be CPGs (**Fig. 4c** and **Supplementary Fig. 4b-d**). First, 9.9% of OMIM genes (*N* = 30), as well as three CPGs (*BRCA1/2* and *XPA*), exhibited a strong prevalence of germline variants in cancer compared to controls as well as high frequent biallelic inactivation. We refer to these as ‘Canonical CPG-like type1’ OMIM genes. The most representative OMIM genes in this cluster were *LRRC6*, *GSS*, and *SRP72* (**Fig. 4c**-**e** and **Supplementary Table 5**). A second cluster showed prevalence of germline variants in cancers but did not preferentially follow a ‘two-hit’ model. This cluster included six CPGs (*APC*, *ERCC3*, *FANCA*, *FANCG*, *MPL,* and *WRN*), named as CPGs with ‘low biallelic inactivation’. 48.4% of OMIM genes (*N*=147) were assigned to the second cluster including *EIF3F*, *SGSH*, and *ACADM*. The third cluster of genes showed high tissue-specificity with relatively low two-hit preferences including three CPGs (*BLM*, *BUB1B,* and *SERPINA1*), named ‘High tissue-specific expression’. This class included 26.0% of OMIM genes (*N*=79) with *SERPIND1*, *TEX11*, and *PAH* which showed exceeded expression in a specific tissue such as the liver or kidney. The fourth cluster implicates another ‘broadly expressed CPG-like’ gene set which showed a high two-hit preference with broad expression including three CPGs (*ATM, CHEK2,* and *FANCC*) with 8.2% of OMIM genes (*N*=25) with *SMPD1*, *AK7*, and *KPTN* (**Fig. 4c-e** and **Supplementary Fig. 5**). The last cluster also showed strong ‘two-hit’ preferences with more specific expression and included two CPGs (*RECQL4* and *TP53)* with 7.6% OMIM genes (*N*=23). This cluster was therefore named the ‘Canonical CPGs with strong two-hit preference’ cluster (**Fig. 4c-d** and **Supplementary Fig. 4c**-**d**).

### *PAH* is a potential cancer-predisposing gene

For a deeper understanding of the role of OMIM genes as possible new CPGs, we focused on phenylalanine hydroxylase (*PAH*). *PAH* plays a key metabolic role in converting the L-phenylalanine to L-tyrosine. Inherited variants in the *PAH* gene increase the conversion of L-phenylalanine to L-phenylpyruvate, a metabolic feature of phenylketonuria (PKU), which is an autosomal recessive genetic disease with symptoms that include intellectual disability, seizures, nausea, and vomiting ^23^. It showed the highest frequencies of pathogenic germline variants (Tier 1 and 2) in a Pan-cancer in both the PCAWG and TCGA datasets (2.2% in PCAWG, 2.1% in TCGA compared to 0.7% in KG; **Fig. 4a** and **Supplementary Fig. 4a**). *PAH* was grouped in Cluster 3 which shows strong germline prevalence in a Pan-cancer with liver-specific gene expression (**Supplementary Table 5**).

To evaluate metabolic changes associated with *PAH* variants, we conducted metabolome analysis for plasma samples from 8 *PAH* heterozygote carriers and 22 healthy donors (i.e., non-*PAH* carriers; see **Methods** and **Fig. 5a**). As a result, we identified 40 metabolites with significantly different concentrations, including 8 up-regulated and 32 down-regulated metabolites, in *PAH* heterozygote carriers compared to healthy donors (*P* < 0.05 by Welch’s T-test, **Supplementary Table 6**). As expected, we observed increased L-phenylalanine levels (Phe; 1.3-fold, P = 6.39 × 10−4) in the blood of *PAH* carriers, a well-known characteristic of *PAH* deficiency (**Fig. 5a**). Intriguingly, PCA analysis for the 220 measured metabolites, including 40 metabolites with significantly different concentrations, clearly distinguished *PAH* carriers and non-carriers, indicating that metabolite levels indeed appeared to be affected by the expression of *PAH* variants (**Fig. 5a**). To better understand the metabolic effects of *PAH* variants, metabolic simulations were conducted by using patient-specific genome-scale metabolic models (GEMs) ^24^ (**Supplementary Fig. 6a**). Input data (i.e., metabolite uptake rates) were obtained from simulation of GEMs presenting 3 *PAH* carriers and 309 *PAH* non-carriers in the PCAWG-TCGA Liver-HCC samples as a model (see **Methods**). The GEMs predicted that L-phenylalanine was more actively secreted in the *PAH* carriers than the *PAH* non-carriers (*P* = 1.42 × 10^−3^ by Student’s t-test), and L-phenylalanine was more actively converted to L-tyrosine in the *PAH* non-carriers. Furthermore, we conducted disease-based enrichment analysis by using 110 target metabolites (see **Methods**). As a result, the *PAH* dysfunction group appeared to be the most significantly related to inflammatory disease (*P* = 1.55 × 10^−8^, **Supplementary Fig. 6b**), followed by diseases related to the heart muscle or optic atrophy (*P* = 1.55 × 10^−8^, and *P* = 5.26 × 10^−8^, respectively). This indicates that metabolic changes occurring in the *PAH* germline variants may also affect multiple diseases, especially those associated with immune system dysfunction, in addition to PKU.

**Figure 5.**
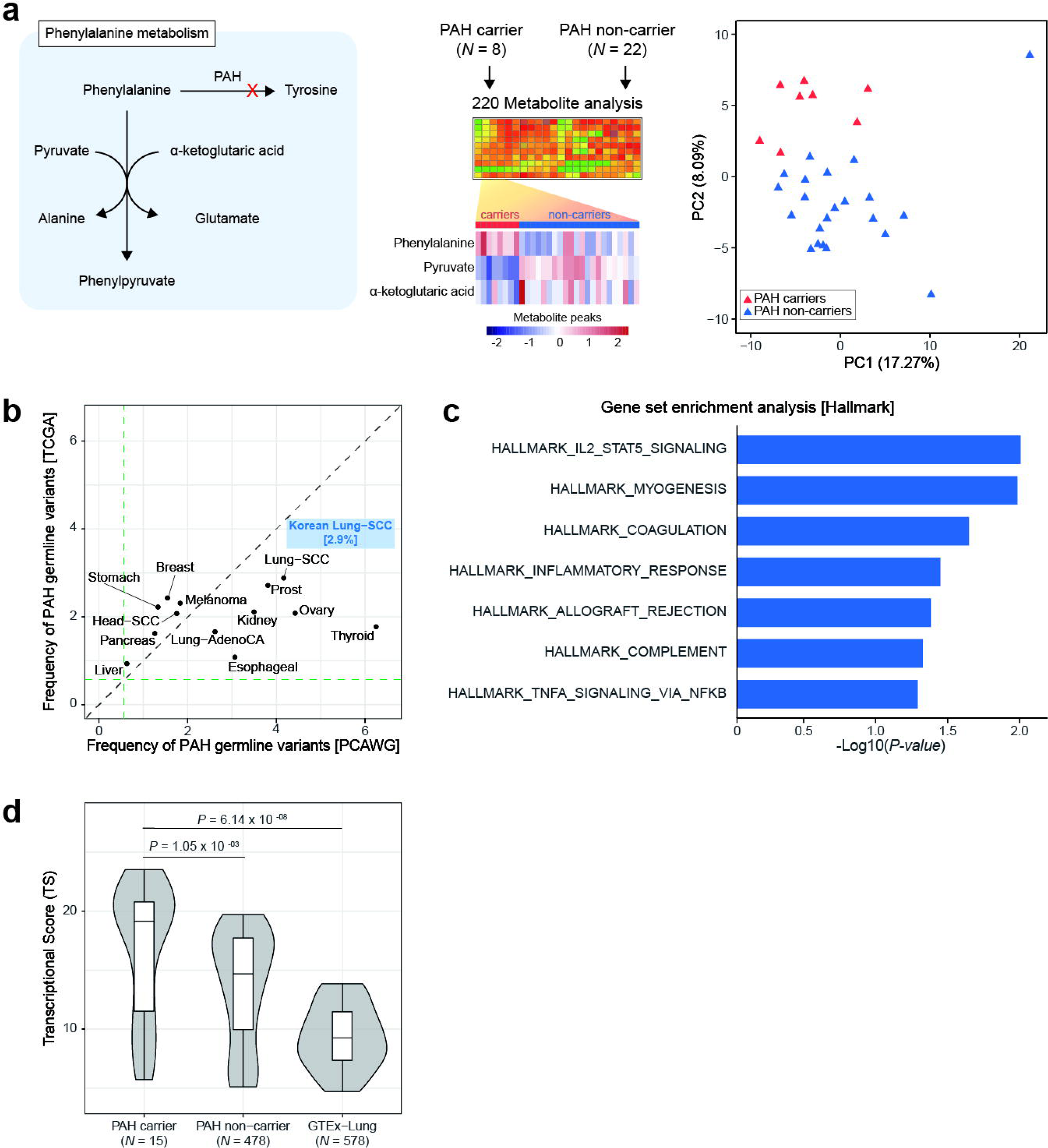
Analysis of the metabolic effects of *PAH* variants. (a) Diagram of phenylalanine hydroxylase (*PAH*) pathway and profiling of 220 metabolites for the *PAH* carriers and *PAH* non-carriers. In phenylalanine metabolism, L-phenylpyruvate can be generated from L- phenylalanine via two enzymatic reactions, one with pyruvate and the other one with α-ketoglutarate. Three representative metabolites are presented in the heatmap (red: higher metabolite peaks in *PAH* carriers, blue: lower metabolite peaks in *PAH* carriers). PCA using the 220 metabolites clearly distinguished the *PAH* carriers from the *PAH* non-carriers. **(b)** The frequencies of *PAH* pathogenic germline variant across cancer types in PCAWG (x-axis) and TCGA (y-axis; after removing overlapping samples with PCAWG) datasets. The frequency of *PAH* in an independent cohort, Korea Lung cancer carcinoma, is presented in blue. Green dashed lines represent the PAH frequency in 1KG (0.7%). **(c)** Hallmark GSEA signatures from PCAWG/TCGA lung cancer RNA-sequencing data are ranked by normalized enrichment score (NES), which represents the enrichment of each cancer hallmark in PAH carriers compared to non-carrier samples. Hallmark signatures are ranked by P-values (at nominal P <0.05). (**d)** Box plot showing the transcriptional score (TS) using immune checkpoint-related gene set (*N*=38) across three groups: *PAH* carrier in Lung-SCC PCAWG/TCGA, *PAH* non-carrier in Lung-SCC PCAWG/TCGA, and normal lung samples from GTEx. High TS indicates increased immune microenvironment correlation within a group.

Next, to understand the detailed tumorigenic mechanism of *PAH*, we selected Lung-SCC which shows the top-ranked frequencies of pathogenic germline variants in both PCAWG (4.2%) and TCGA (2.9%) across cancer types (**Fig. 5b**). We confirmed the lung-specific prevalence of *PAH* pathogenic germline variants using independent whole-exome sequencing (WES) data of Korean Lung-SCC samples (2.9%; 7 out of 245, **Fig. 5b**). However, the position of pathogenic germline variants between PCAWG-TCGA and Korean Lung-SCC patients were different (**Supplementary Fig. 6c**). To test whether lung-specific oncogenic processes differ with *PAH* state, we performed a gene set enrichment analysis (GSEA) using genes which are expressed in both *PAH* carriers and non-carriers in Lung-SCC from PCAWG-TCGA (see **Methods**). Interestingly, immune-response related cancer hallmarks were strongly enriched in *PAH* carriers such as interleukin-2 and STAT5 signalling, inflammatory response, coagulation, and TNF-α signalling via NF-k (**Fig. 5c**). Furthermore, we measured the transcriptional score (TS), a gene-wide measurement that sums up all the gene expression correlation coefficients between 38 immune checkpoint modulatory genes within a group (see **Methods**) ^25^, across three groups: (i) *PAH* carrier in PCAWG-TCGA Lung-SCC, (ii) *PAH* non-carrier in PCAWG-TCGA Lung-SCC, and (iii) normal lung samples from GTEx ^26^. *PAH* pathogenic germline variant carriers (*N* = 15, median TS = 19.14) showed significantly increased TS compared to *PAH* non-carrier (*N* = 478; median TS = 14.69, *P* = 1.05 × 10^−3^ by Wilcoxon rank sum test) and normal lung samples from GTEx (*N*= 578; median TS = 9.25, *P* = 6.14 × 10^−8^ by Wilcoxon rank sum test, **Fig. 5d**). This indicates that the gene expression levels of immune check-point modulatory genes in *PAH* carriers tend to positively correlate with each other, similar to a hot tumor microenvironment, which shows high response rates to immune checkpoint inhibitors ^27^. Additionally, we checked the mutational signatures present based on the somatic mutation patterns, but we did not observe any differences in the type of signatures or frequencies of each signature between the *PAH* carrier and non-carrier group (**Supplementary Fig. 6d**). Altogether our data validate that *PAH* may be a candidate of CPGs which modulates immune-associated pathways to contribute to cancer.

## Discussion

Here we found that rare pathogenic variants in some OMIM genes contribute to cancer risk by analyzing 2,642 tumor samples. Specifically, OMIM genes in metabolic and immunological disease classes showed a substantial enrichment of pathogenic germline variants in cancer compared to controls (median Log2OR = 0.35, *P* = 7.35 × 10^−2^ and median Log2OR = 0.47, *P* = 2.91 × 10^−2^, respectively). Our study further reveals five possible clusters of OMIM genes by integrating four different complementary features (**Fig. 6**). In our study, we uncover a diverse range of CPGs from OMIM genes, including new candidates linked to cancer susceptibility.

**Figure 6.**
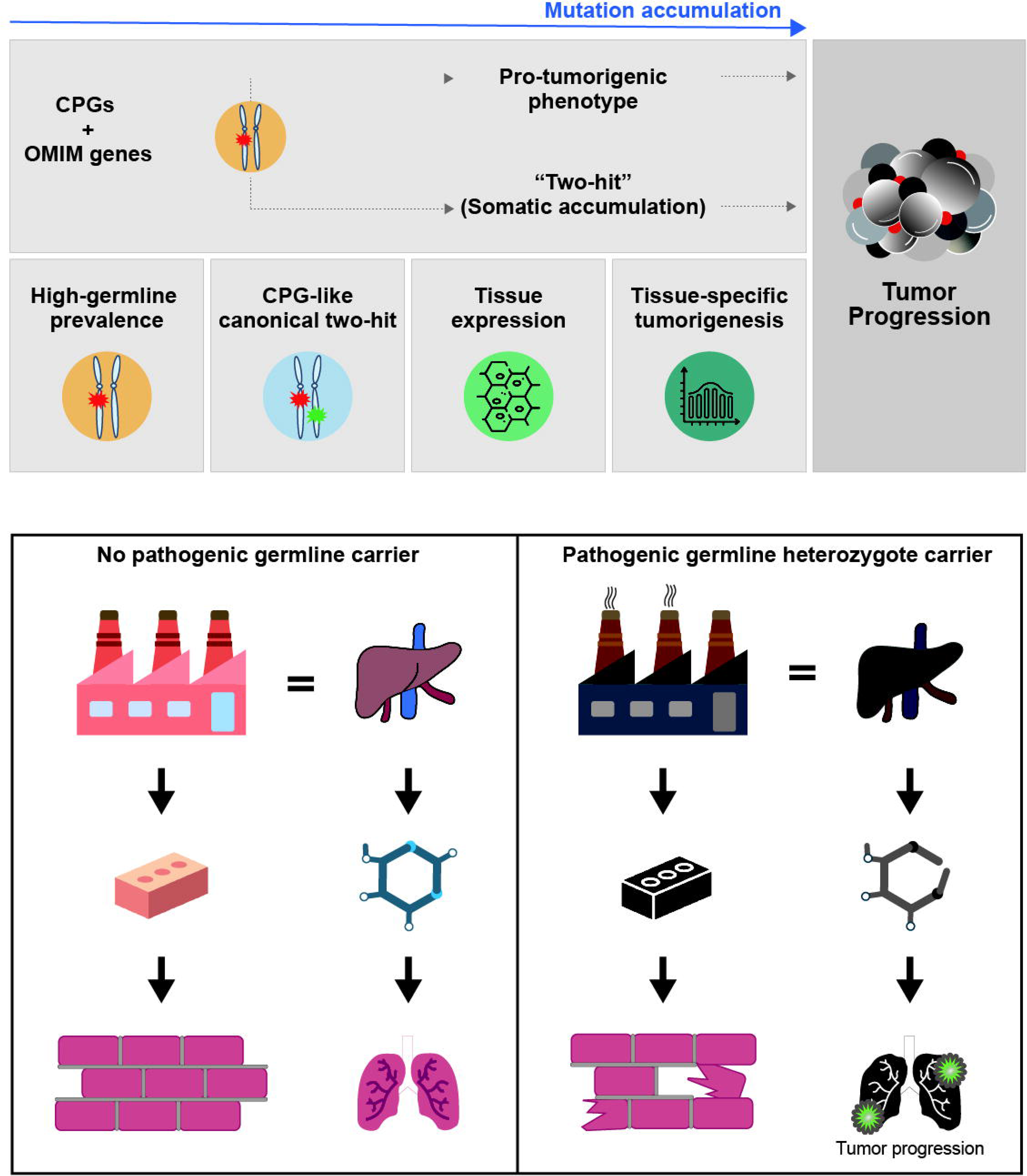
The impact of hereditary germline variants on tumorigenesis. Summary describing our hypothesis that rare germline variants are linked to increased cancer risk at a level similar to previously known CPGs (top panel). Possible tumorigenesis mechanisms along with abnormal development following pathogenic germline variants are shown in the bottom panel. These OMIM genes were classified using four features that comprise germline variant enrichment, two-hit preference, gene expression across tissue types, and tissue- specificity.

First, our case-control analysis demonstrates that rare germline variants in Mendelian disease-associated genes may increase the lifetime risk of developing cancer similar to known CPGs. Of note, disease class-based enrichment analysis showed an increased cancer risk with genes involving immune, endocrine, and metabolic disorders. In fact, immuno-oncology ^28, 29^ and onco-metabolism ^5, 30, 31^ have become important areas of oncology research during the last decade. However, only a few germline variants linked to immune-oncology and cancer-metabolism are current targets for therapeutic development ^32, 33^. We believe our observations are closely related to these important hallmarks of cancer and will provide leads for novel treatment strategies.

We observed enrichment of rare germline variants in autosomal recessive OMIM genes by case-control analysis. Genetically, this might be related to the possible diverse clinical phenotype of heterozygote rare germline variant carriers. The common expectation for rare variant heterozygotes is that carriers have a normal lifespan since the wild-type allele maintains gene function. However, prolonged life expectancy may allow unexpected phenotypes attributable to rare germline variants to emerge in heterozygotes. For example, heterozygote carriers may have an altered metabolite profile that does not provoke a classic disease phenotype. However, long-term exposure to this altered metabolism may negatively impact health and physiology. There may also be gene-environment interactions in heterozygous carriers of OMIM genes where a subtle environmental change can trigger cancer.

In the 40 years since Alfred Knudson’s ‘two-hit’ hypothesis, many classical cancer predisposition genes have been identified and validated in particular tissues or across cancers^2^. Our study suggests that the ‘two-hit’ hypothesis still remains a useful model for uncovering and classifying cancer genes, but also exceptions where pathogenic germline variants are not coupled with additional somatic alterations, even in CPGs. Genes in Cluster 3 show significantly high tissue-specific gene expression compared to other clusters (mean tissue-specificity value in Cluster 3 = 14.13, other cluster = 2.21, *P* < 2.2× 10^−1^^6^ by Student’s t-test), providing evidence for a non-classical mechanism where bi-allelic inactivation is not commonly observed in a Pan-cancer. Accordingly, we suggest that an indirect carcinogenic mechanism would be related to emerging hallmarks of cancer such as cell senescence, non-mutational epigenetic reprogramming, and unlocking phenotypic plasticity ^5^.

In this study, we present *PAH* as a novel candidate CPG that carries frequent germline variants and is assigned to Cluster 3. Interestingly, although *PAH* is scarcely expressed in lung epithelium, the enrichment of *PAH* germline variants was strongly observed in Lung-SCC . Transcriptomic analysis revealed that squamous cell lung cancer patients with *PAH* rare variants were associated with the immune-related pathways that could be influenced by long-lasting dysregulated metabolic stress. Indeed, metabolome data as well as patient-specific GEMs from *PAH* carriers showed metabolic profiles that were clearly distinct from the normal population. Our finding provides an insight to understand the possible mechanism of cancer progression through pathogenic germline variants in *PAH* since comorbidity between phenylketonuria and multiple cancer types has been observed ^34, 35^.

The sequencing of an even larger number of tumors will further refine the clustering and will also allow a complete description of the classes that differently contribute to cancer. One possible caveat is that a similar power limitation is caused by population biases since PCAWG is mainly derived from European/American patients (72.1%). Some population-specific pathogenic variants – especially non-European variants – could be underestimated in our study due to the limited sample size. The sequencing of an even larger number of tumours and healthy control individuals will allow further refinement as we validated *PAH* enrichment with Korean lung cancer data and will also provide a clearer picture of the diversity of OMIM genes that contribute to cancer through different mechanisms.

## Methods

### Ethical approval

This paper reanalyzes previously published data sets. All cancer patient and healthy controls data were handled in accordance with the policies and procedures of the Seoul National University Hospital.

### Tumor sequencing data from PCAWG

The final consensus set of merged germline mutation calls in variant call format (VCF) generated by the Pan-cancer Analysis of Whole Genomes (PCAWG) ^14^ consortium was obtained from the International Cancer Genome Consortium Portal (https://dcc.icgc.org/releases/PCAWG) with authorization. The final dataset contains 2,642 high-quality samples after excluding 192 samples with possible technical issues from 2,834 donors ^14^. The germline variant calling file was separated into two germline VCF files: one containing 1,823 donors from the non-US International Cancer Genome Consortium (ICGC) and the other containing 819 donors from The Cancer Genome Atlas (TCGA). Separate VCF files were integrated using the bcftools-1.9 as recommended by the ICGC data coordination center (DCC) ^36^. The final set of PCAWG germline calls comprises 2,642 donors who are composed of four subpopulations as described in **Supplementary Fig. 1c** and **Supplementary Table 1**: European/American (1,904 donors; 72.1%), East Asian (396 donors; 15.0%), African (125 donors; 4.7%), and South Asian (36 donors; 1.4%). This also includes 181 donors (6.9%) whose ethnic information was not identified. The germline variant calling was originally performed by the ICGC working group ^14^ assembling the SNVs, indels, and structural variations (SV) using the six different variant callers; GATK HaplotypeCaller, FreeBayes, Real Time Genomics (RTG), Delly, TraFiC mobile element insertion caller (https://gitlab.com/mobilegenomes/TraFiC), and Eagle2.

### Control exomes from 1000 Genomes Project

Exome sequences of healthy controls were collected from 1000 Genomes Project (1KG; Phase III high-coverage whole-exome sequences) ^15^ VCF from the ftp server (ftp://ftp.1000genomes.ebi.ac.uk/vol1/ftp/release/20130502/) under the following authorization (dbGaP phs000710.v1.p1). The data contains information from 2,504 individuals generated from four subpopulations (European/American, *N*=850; African, *N*=661; East Asian, *N*=504; South Asian, *N*=489, **Supplementary Fig. 1c**). Data generation and variant calling procedures are detailed in the 1KG flagship paper ^15^.

### Clinical information

Clinical and histological annotation information for PCAWG donors was obtained from the ICGC Portal ^14^ (https://dcc.icgc.org/releases/PCAWG). PCAWG data included 1,462 males (55.3%) and 1,180 females (44.7%). The Pan-cancer cohort comprises 37 distinct tumor types (**Supplementary Table 1**) and the germline samples were obtained from non-tumor samples mainly from blood but also tissue adjacent to the primary site or other sites such as bone marrow, lymph node. The demographic information for 1KG control samples is available from the 1000 Genome portal (https://www.internationalgenome.org/data).

### Variant annotation and filtering step

To generate a consistent functional annotation set, we compiled each VCF file from PCAWG (*N*=2,642) and 1KG (*N*=2,504). First, variants in ENCODE/DUKE ^37^ and DAC blacklist ^38^ regions were discarded to filter out variants in low mappability regions and only selected variants in the ENCODE/CRG GEM mappability region ^39^ (75mers). Next, we annotated the collected variants in the VCF file using ANNOVAR (version 2014 Apr 14) ^40^. Of the data Annovar reports with filter-based functional annotation, we used (i) the clinical and phenotypic effect of variants via ClinVar ^13^ (accessed on 18 June 2019), (ii) minor allele frequencies of variants across eight populations: African/African American (AFR), South Asian (SAS), East Asian (EAS), Latino/Admixed American (AMR), non-Finnish European (NFE), Finnish European (FIN), Ashkenazi Jewish (ASJ) and Other (OTH) and also globally in The Genome Aggregation Database (gnomAD) exome (v2.1.1) ^12^. Simultaneously, Variant Effect Predictor (VEP)-release-96 ^41^ was used to explicate the gene-based information of canonical transcripts. Of the data VEP reports, we used the consequences of the protein-coding variants: synonymous, missense, stop gain and loss, splice site, frameshift indel, and in-frame indel. Next, we discarded all variants marked as possible technical artifacts in the gnomAD exome including excess heterozygosity at a variant site (InbreedingCoeff), allele count zero variants after filtering out low-confidence genotypes (AC0), and failed random forest filtering threshold (RF).

### Principal component analysis using common variants

Many germline variants from whole-exome sequencing data or genome-wide association studies are expected to vary according to different ethnicities present within the cohort. To check the possibility of the potential confounding effects from population stratifications, we performed a PCA with only the common germline variants (non-synonymous) with ≥ 5% MAF from both globally in PCAWG and 1KG and also each ethnic group of PCAWG/1KG (four subpopulations) and 90% genotyping rate using PLINK ^42^ version 2.0.

### Rare pathogenic germline variants

Rare variants were defined as those whose frequency was < 0.5% globally and less than 1% in each of the eight subpopulations in gnomAD exome. For variants that are not present in the gnomAD exome, we checked the number of detections in our cohorts (PCAWG, 1KG separately) across variants to remove possible technical artifact variants, and discarded variants where more than 1% samples were detected in either PCAWG or 1KG cohort (9.4% of variants; 139,678 out of 1,485,345).

From collected rare variants, pathogenic variants were defined as two conditions: (1) Tier1 variants (potentially deleterious) were defined as premature protein-truncating variants (PTVs) including splice donor/acceptor site, frameshift indel, stop gain/lost variant as well as not annotated as Benign or Likely benign in ClinVar ^13^. (2) Tier2 variants were defined as variants with ‘Pathogenic’, ‘Likely pathogenic’, ‘association’, and ‘risk factors’ with related clinical evidence developed from the American College of Medical Genetics and Genomics (ACMG) in ClinVar ^13^ .

### Gene set classification

We retrieved the genes with known disruption resulting in clinical diseases and phenotypes from the Online Mendelian Inheritance in Man (OMIM) databases ^7^ (https://www.omim.org/; acquired Jan 21, 2020) via the Gene Map (*genemap2.txt*) and Morbid Map (*mim2gene.txt*). The OMIM database provides 17,076 disease-associated genes from 5,392 diseases represented in the Gene Map (cytogenetic locations of genes) and Morbid Map (cytogenetic locations of disorders). Next, 5,460 disease-associated genes which are associated with at least one phenotype (genetic disease) were selected from *genemap2.txt* for further analysis. We then cross-checked genes represented in *mim2gene.txt* and annotated them to Ensembl gene name using *annotables* package in R (https://github.com/stephenturner/annotables); finalizing to 4,095 unique genes. Moreover, a total of 152 known germline CPGs were collected from recent studies ^16^ including 114 from a recently published review paper ^2^, 11 from Cancer Gene Census-Germline (http://cancer.sanger.ac.uk/census/), 12 from references search (details in from Huang *et al* ^16^), and 15 genes from the St. Jude PCGP germline study ^43^. For collecting high-confidence somatic driver genes (SODs), we first collected 678 genes across three data sources: IntOGen ^44^, MutSig ^45^, and MutPan ^46^. To assign non-overlapping gene sets, CPGs (*N* = 152) set was firstly defined, and then 3,952 OMIM genes were defined after discarding 143 overlapping genes with CPGs. Thereafter, 257 SODs were collected after removing 421 overlapping genes, either CPGs or OMIM genes. Subsequently, 53 CPGs that have no clinically pathogenic variants found in a TCGA-based germline variant analysis ^16^ were excluded from the final CPG set. Finally, 4,308 genes were defined as input genes including 99 CPGs, 257 SODs, and 3,952 OMIM genes including 2,007 autosomal-recessive [AR] genes, 1,194 autosomal-dominant [AD] genes, 252 AD-AR genes (i.e., variants from the same gene have dual effects either dominant or recessive), 211 X-linked genes, 53 somatic, 3 digenic, 5 isolated, 13 multifactorial, 2 Y-linked and 212 genes with unknown inheritable phenotypes were finally selected based on the distinctive inheritance annotation (**Supplementary Table 2**). As a control set, 5,598 genes were collected from all human genes found in PCAWG and 1KG except for CPGs, SODs, OMIM genes, 196 particular disease/cancer-related genes (https://diseases.jensenlab.org/Downloads), 100 uncharacterized proteins and 126 DNA repair genes from Ensembl gene description annotated using biomaRt ^47^ packages in R.

### Statistical analysis of pathogenic germline variant enrichment

Burden test analysis was performed by comparing the frequency of pathogenic germline variants within a gene between cancer cases (PCAWG) and controls (1KG). This method is designed to measure gene-level cancer risk enrichment by collapsing possibly rare pathogenic germline variants using a generalized linear regression model (GLM) with stats package in R. The model was performed in each gene-tissue (Pan-cancer and single 26 cancer types with > 20 samples) pair as follows:

glm (*N* ∼ *Germline Variants* + *PC1 + PC2*, *family= ’’binomial’’*)

where: *N* = case (1) or control (0), *Germline Variants* = indicating the number of samples that carries rare pathogenic germline variants for each gene-tissue pair, PC values from PCA analysis of PCAWG and 1KG to control population structures were used as an input for the regression. The regression coefficient and *P*-value were computed for individual gene-tissue pairs using the *summary* function in R. More positive coefficient values refer to stronger enrichment of pathogenic germline variants in the case compared to control.

### Pathogenic germline variant enrichment in disease class and pathways

To systematically estimate the enrichment of pathogenic germline variants in disease classes compared to controls, we collected 10 diseases classes (Cardiovascular, Endocrine, Epilepsy, Hematology, Immune, Liver, Lung, Metabolism, Muscular/Skeletal and Neurological) with 96 hereditary diseases based on clinically relevant disease diagnosis from Centogene (https://www.centogene.com/diagnostics/ngspanels.html) and Blueprint Genetics (https://blueprintgenetics.com/tests/panels/). From 3,952 OMIM genes, 784 genes were mapped to the 10 disease classes with 91 OMIM diseases.

From the Molecular Signatures Database ^48^ (MsigDB version 7.0), we collected 186 KEGG signalling pathways. To remove possible redundancy between KEGG pathways, a gene set over-representation analysis was done by incorporating network-based gene weights by using functional Link Enrichment of Gene Ontology or gene sets (LEGO V2.0) ^49^. Finally, 70 KEGG signalling pathways were collected with 1,721 OMIM-associated genes.

The enrichment analysis was performed in each disease class or pathway across cancer types as follows:

glm (*N* ∼ *Pathway or Disease classes* + *PC1* + *PC2*, *family=’’binomial’’*)

Where: *N* = case (1) or control (0), *Pathway or Disease classes* = indicating the number of samples that carry a rare pathogenic germline variant for each disease class or pathway-related genes. PC values from PCA analysis of PCAWG and 1KG were used as an input for the regression model.

### Copy-number alteration from PCAWG

Genomic copy-number alteration (CNA) data in gene-level calls ^50, 51^ across 2,642 samples were also obtained from the ICGC data portal (final_consensus_passonly.snv_mnv_indel.icgc.public.maf and final_consensus_passonly.snv_mnv_indel.tcga.controlled.maf). It provided the summarized intersections of genomic region calls from 6 copy-number alteration callers, ABSOLUTE; ACEseq; Battenberg; cloneHD; JaBbA; and Sclust, run as described in Dentro *et al* ^52^. From the CNA data, copy-number loss of heterozygosity (LOH) was defined as following in the PCAWG paper ^51^ when the minor allele was zero.

### Statistical analysis of two-hit preference analysis

The excess of pathogenic germline variants in samples with LOH compared to samples without an LOH event was performed to test the two-hit hypothesis at the gene level across cancer types. The model was performed in each gene-tissue pair as follows:

glm (*N* ∼ *Germline Variants* + *PC1 + PC2*, *family=’’binomial’’*)

Where: *N* = samples with LOH event (1) or samples without LOH event (0), *Germline Variants* = indicating the number of samples that carry rare pathogenic germline variants for each gene-tissue pair, PC values from PCA analysis of only PCAWG samples to control population structures within a cancer cohort were used as an input for the regression. The regression coefficient and *P*-value were computed for individual gene-tissue pairs using the *summary* function in R. More positive coefficient values refer to stronger enrichment of pathogenic germline variants in samples with LOH compared to samples without LOH.

### PCA for gene classification

To classify OMIM genes and CPGs, we focused on genes with 1.5-fold more frequent rare germline variants in both PCAWG and TCGA compared to the controls (1KG). Then, we integrated four features including (i) germline variant prevalence in cancer cohorts compared to healthy control (case-control analysis by regression model; Log2 odds ratio), (ii) two-hit preferences (excess of pathogenic germline variants in samples with LOH event compared to samples without LOH event by regression model; Log2 odds ratio), (iii) number of tissues with expression level higher than the mean expression value of each gene across 20 tissue types from Human Protein Atlas (HPA), (iv) tissue-specificity which indicates the expression level of genes are broadly expressed or specifically expressed across cancer types (**Fig. 4b**). In HPA, normalised expression level (NX) greater than 2 has been chosen to define a tissue-specific high expression after checking the distribution of expression levels across genes. We applied these four values as an input for PCA based clustering analysis using *prcomp* function in stats R package. We used the first three PCs to cluster the individual input genes and clustered genes into four clusters including outliers using the tclust package in R (https://cran.r-project.org/web/packages/tclust/index.html) ^53^ (**Supplementary Fig. 4b**).

### Independent cancer cohort from TCGA

For the independent validation, TCGA germline variants for 10,389 samples ^16^ (PCA.r1.TCGAbarcode.merge.tnSwapCorrected.10389.vcf) was downloaded from the National Cancer Institute (NCI) Genomic Data Commons (GDC) ^54^ legacy archive (dbGaP phs000178). It contains 7,573 European (72.9%), 901 African American (8.7%), 651 Asian (6.3%), and 1,264 other or unknown ethnicity (12.2%). After excluding 782 samples which are overlapped with the PCAWG study for further analysis, 9,607 TCGA samples were finally collected.

### Metabolome analysis of *PAH* carriers

We also collected blood samples from the parents of 8 Phenylketonuria patients to obtain data from *PAH* heterozygote carriers and also 22 healthy controls who do not have *PAH* pathogenic germline variants. The samples were prepared according to the procedure and policies of the Seoul National University Hospital [IRB No. H-2101-196-1193] and transferred to Human Metabolome Technologies, Incorporation (HMT) to measure the metabolic changes with *PAH* heterozygote carrier. The samples were treated according to the protocol provided by HMT (Document ID: BLB.1.0.0). The metabolic changes were evaluated using the Cation and Anion modes of CE-TOFMS (Capillary Electrophoresis Time-of-Flight Mass Spectrometry) based metabolome profiling. Peaks retrieved from the CE-TOFMS analysis were extracted using automatic integration software (MasterHands ver. 2.18.9.1 created at Keio University), which included m/z, migration time (MT), and peak area.

The following equation was used to convert the peak area to the relative peak area.

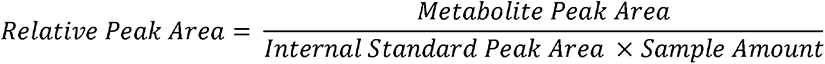

The m/z and MT were then used to assign putative metabolites from HMT’s standard library and Known-Unknown peak library. All metabolite concentrations were measured using standard curves generated from single-point (100µm) calibrations and normalizing the peak area of each metabolite with respect to the area of the internal standard. These quantified concentrations from 110 target metabolites were then used for disease-related enrichment analysis by MetaboAnalyst 5.0 ^55^ (https://www.metaboanalyst.ca/), a web-based metabolome data analysis tool.

### Metabolic simulations using genome-scale metabolic models

Genome-scale metabolic model (GEM) ^24^ was used for metabolic simulations on PAH carriers and PAH non-carriers. GEM is a computational model that contains information on biochemical reactions for entire metabolic genes in a cell, and can be simulated by using optimization techniques for various metabolic studies ^56^. In this study, a previously developed generic human GEM Recon 2M.2 ^24^ was transformed into a context-specific (or patient-specific) GEM through integration with RNA-Seq data from PCAWG-TCGA Liver-HCC and Lung-SCC samples. Task-driven Integrative Network Inference for Tissues (tINIT) method, along with a rank-based weight function, was used to generate patient-specific GEMs. Using the Liver-HCC GEMs, metabolites secreted from the PAH carriers (*N* = 3) and the PAH non-carriers (*N* = 309) were first predicted. As a result, the Liver-HCC GEMs presenting the PAH carriers and the PAH non-carriers are predicted to secrete 16 and 40 metabolites, respectively. The predicted metabolite secretion rates from the PAH carriers with Liver-HCC served as input for the GEMs that present the PAH carriers having Lung-SCC (*N* = 15). Likewise, the predicted metabolite secretion rates from the PAH non-carriers with Liver-HCC were used as input for the GEMs presenting the PAH non-carriers with Lung-SCC (*N* = 478). The maximum uptake rate for the predicted metabolites from the Liver-HCC GEMs was arbitrarily limited to 1.1 times the average metabolite secretion rates in order to simulate the Lung-SCC GEMs. Exchange reactions for inorganic nutrients were set to have the maximum secretion and uptake rates of 1,000 and -1,000 mmol/gDCW/h, respectively. Uptakes of all the other nutrients were not allowed when simulating the Lung-SCC GEMs. Metabolic fluxes of the patient-specific GEMs were predicted by using the least absolute deviation (LAD) method that attempts to minimize the distance between transcript expression level from RNA-seq data and target reaction fluxes to be calculated. tINIT with the rank-based weight function as well as LAD method was implemented using in-house Python scripts ^24^.

### Functional analysis with *PAH*

For the *PAH* analysis in Lung-SCC, we integrated 493 samples from PCAWG and TCGA: 48 from PCAWG (all were derived from TCGA) and 445 from TCGA (not included in PCAWG). RNA sequencing (RNA-seq) data for Lung-SCC samples, were obtained from GDC Data portal (https://portal.gdc.cancer.gov) and calculated the gene expression value after transcriptomic reads alignment to the human reference genome (hg19) using spliced transcripts alignment to a reference (STAR) version 2.5.3a ^57^ and RSEM v1.3.0 (RNA-seq by Expectation Maximization) ^58^. GSEA ^59^ was performed using the java GSEA application version 4.1.0 to evaluate the functional enrichment in Cancer Hallmark with 1,000 permutations. Furthermore, we calculated the transcriptional score (TS) ^25^ which is defined as an absolute number of summed gene expression correlation coefficient values using 38 immune checkpoint modulatory genes ^27^. We compared TS values among three groups of samples (1) *PAH* carrier in Lung-SCC, (2) *PAH* non-carrier in Lung-SCC, and (3) normal lung samples in GTEx ^26^. Since the number of *PAH* carriers in Lung-SCC is much smaller than either *PAH* non-carrier in Lung-SCC or GTEx lung normal sample, the same number of samples were randomly selected 1,000 times (N = 45 samples of non-*PAH* and GTEx normal lung group, separately). The mean TS values were calculated from random sampling with non-*PAH* and GTEx samples were compared with TS value of *PAH* carriers.

To compare somatic mutational signature differences between *PAH* carriers and non-carriers in Lung-SCC, we downloaded the catalog of mutational signatures to the pattern of Single Base Substitutions (SBS) from sigProfiler ^60^ across multiple major cancer types included in the PCAWG and TCGA (PCAWG_sigProfiler_SBS_signatures_in_samples.csv, and TCGA_WES_sigProfiler_SBS_signatures_in_samples.csv, respectively which are available at https://www.synapse.org/#!Synapse:syn11726601/wiki/513478).

### Korean Lung-SCC germline variant calling

For the independent validation of pathogenic germline variants enrichment of *PAH* in lung cancer, we collected 245 peripheral blood mononuclear cell (PBMC) specimens from Korean lung squamous cell carcinoma with informed consent from Samsung Medical Center (SMC) [IRB No. 2013-10-112 and 2008-06-033]. They were processed with the Swift 2S Turbo DNA library kit and whole exome sequencing (WES) data was generated with the IDT xGen Exome Research Panel v1.0 kit on Illumina NovaSeq 6000 platform. The variant calling step was performed using Genome Analysis Toolkit (GATK) Best Practices Workflow pipeline such as HaplotypeCaller ^61^ and we performed variant annotation following the same process described above for PCAWG and TCGA.

## Acknowledgements

This research was supported by a grant of the Korea Health Technology R&D Project through the Korea Health Industry Development Institute (KHIDI), funded by the Ministry of Health & Welfare, Republic of Korea (grant number: HI18C1876) and the National Research Foundation of Korea (NRF; grant number: 2021R1A2C3005360). This work was also supported by the Seoul National University Hospital Research Fund (grant number: 1120190020 and 03-2020-0380) and the computing resources by Global Science experimental Data Hub Center (GSDC) and Korea Research Environment Open NETwork (KREONET). S.P. is supported by the Agencia Estatal de Investigación, Ministerio de Ciencia e Innovación (MCIN/AEI/10.13039/501100011033) through the RETOS project PID2019-109571RA-I00 and the Severo Ochoa Centres of Excellence program to the CNIO (MCIN/AEI/10.13039/50110001103).

## Author contributions

S.S., Y.K., and S.P. designed analyses, evaluated the results, and wrote the manuscript. S.S. and S.P. compiled and performed quality control on the exome sequencing data. S.K. performed GSEA and TS comparison depending on *PAH* state. S.L and J.K. provided samples for *PAH*-associated analysis. S.M.L. and H.K. designed and evaluated the metabolic analysis. S.Y. and Y.K. supervised this study. All authors read and approved the final manuscript.

## Declaration of interests

The authors declare no competing interests.

## Data availability

This paper re-analyses PCAWG whole genome sequencing (retrieved from http://dcc.icgc.org/pcawg/), TCGA whole exome sequencing (https://cghub.ucsc.edu/) and 1000 genomes (http://www.internationalgenome.org/). All data sets are available upon request from the ICGC Data Access Compliance Office (DACO; http://icgc.org/daco), TCGA Data Access Committee (DAC) via dbGaP and the study authorization for 1,000 genomes. The data set of rare variants is from the gnomAD Browser version 2.1.1 (https://gnomad.broadinstitute.org/).

## Code availability

Source code is available at https://github.com/sosug3232/YLAB and https://doi.org/10.5281/zenodo.6791873.

---

**Supplementary Figure 1.**
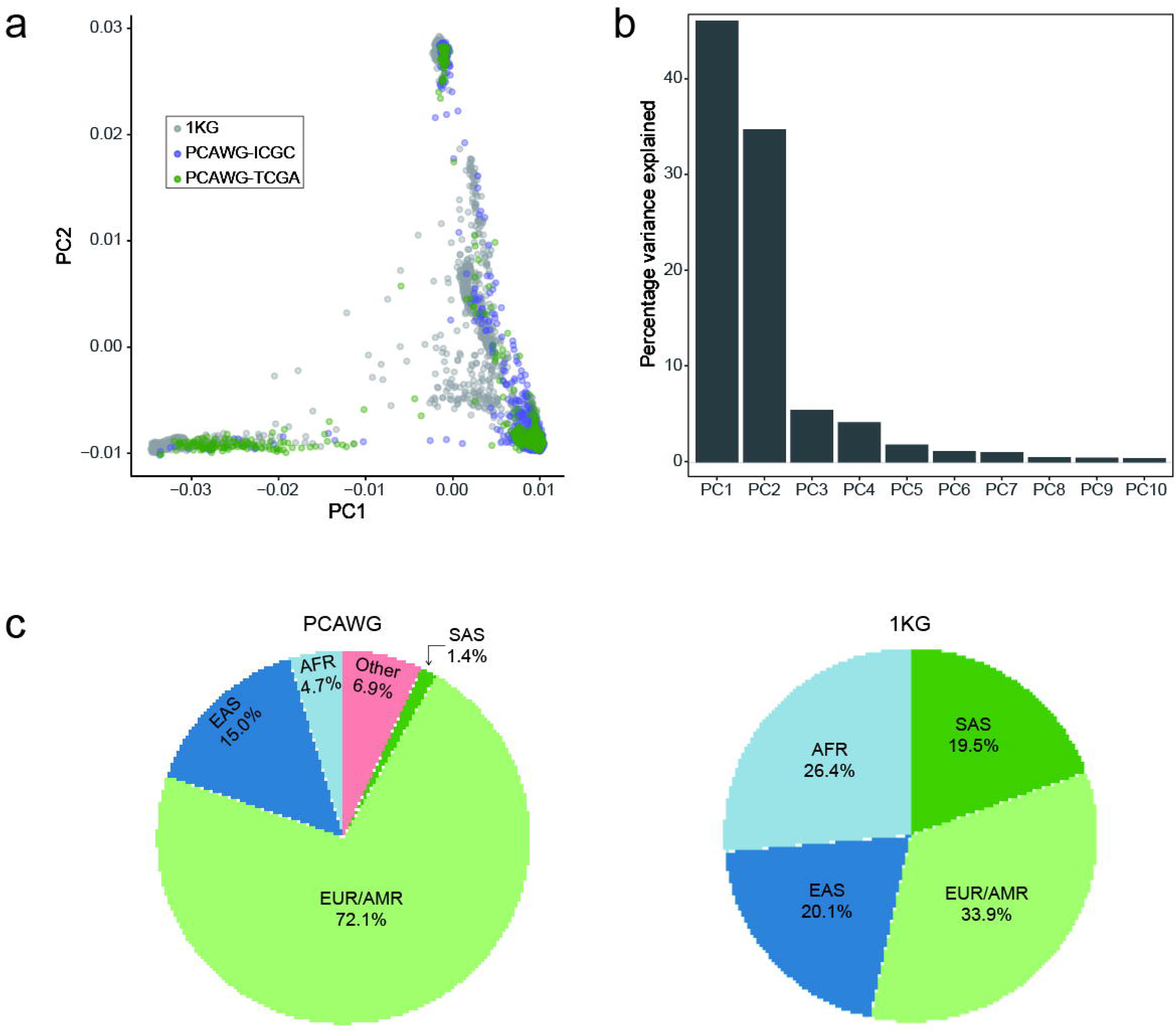
Distribution of population composition and principal components analysis (PCA) using germline variants in PCAWG and 1000 Genomes (1KG). (a) PCA analysis is colored by the source of cohort (blue: PCAWG-ICGC, green: PCAWG-TCGA, and grey: 1KG, respectively). TCGA indicates cancer patients who are in PCAWG that also overlap with TCGA. (b) The percentage of variance is explained for each component after PCA. The first 2 components account for approximately 90°/o of the total variance of the solution. (c) Population composition in PCAWG and 1KG. AFR: African, AMR: American, EAS: East Asian, EUR: European, and SAS: South Asian.

**Supplementary Figure 2.**
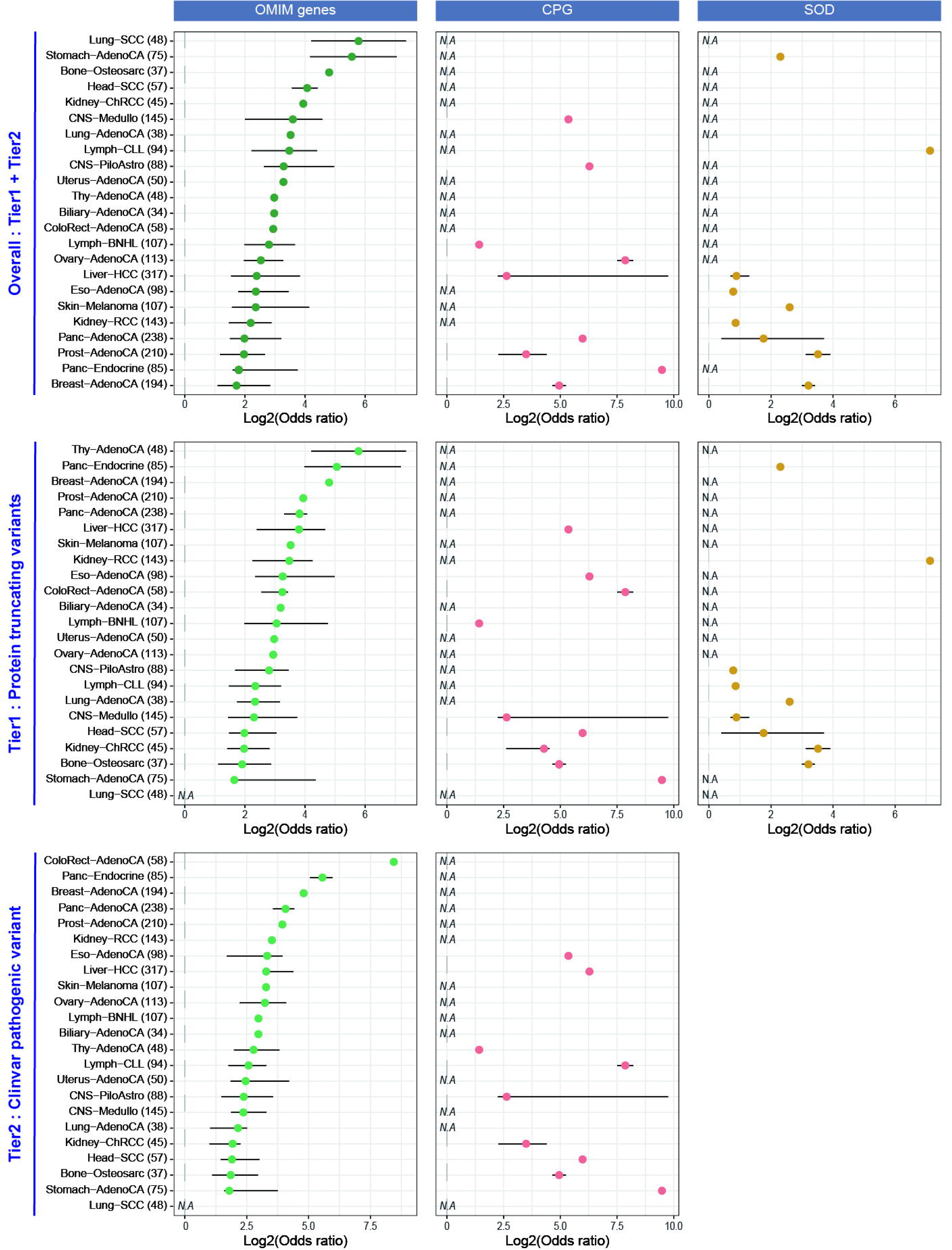
Single-cancer type based case-control analysis across three types of pathogenic variants. Log2 odds ratio distribution with case-control analysis across single cancer types. The length of each whisker is 1.5 times the interquartile range (shown as the height of each box). SODs did not have any Tier 2 variants.

**Supplementary Figure 3.**
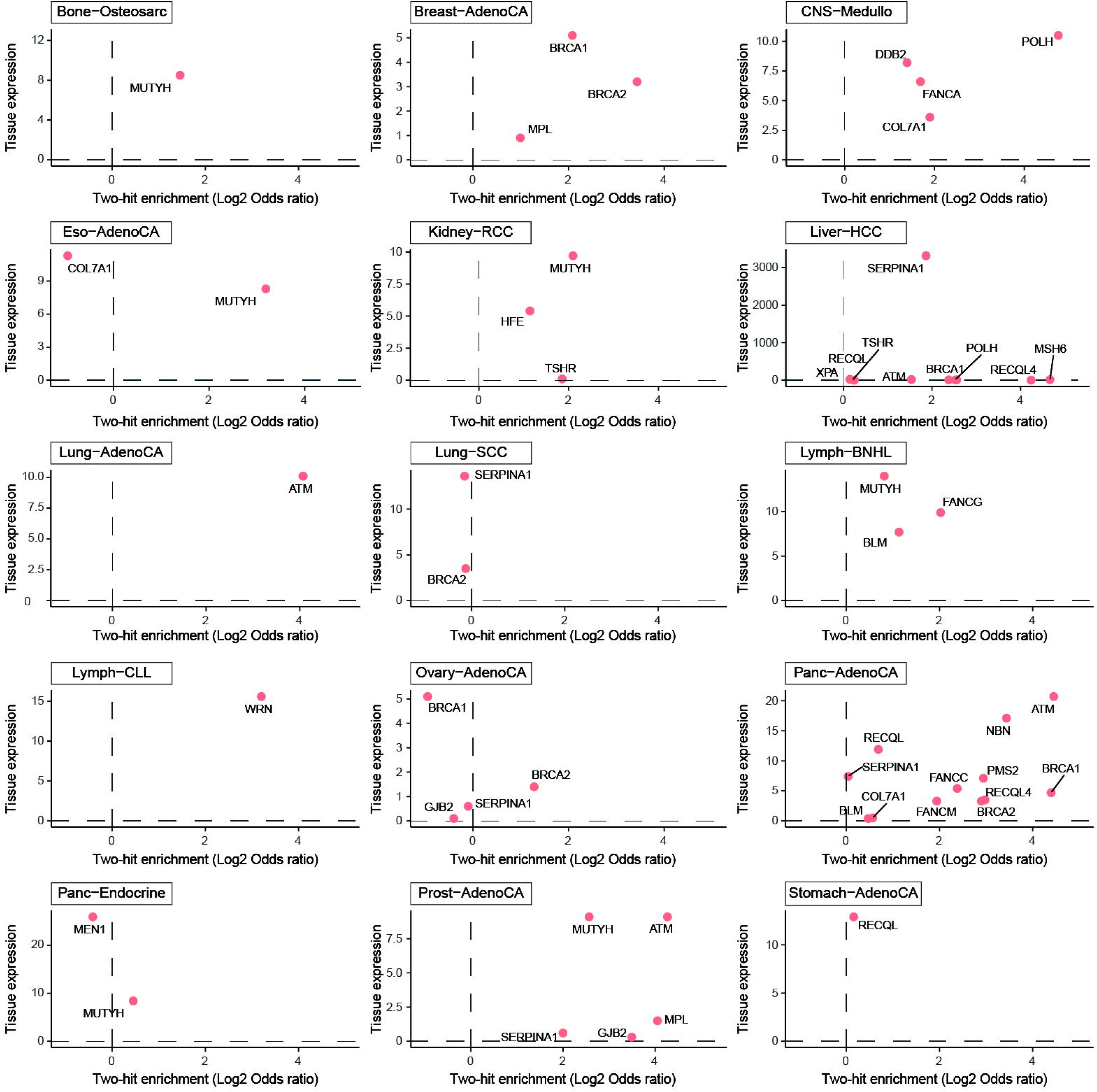
Comparison between gene expression level and two-hit preferences across 15 cancer types. 26 CPGs with at least one pathogenic germline carriers in each cancer types are only selected.

**Supplementary Figure 4.**
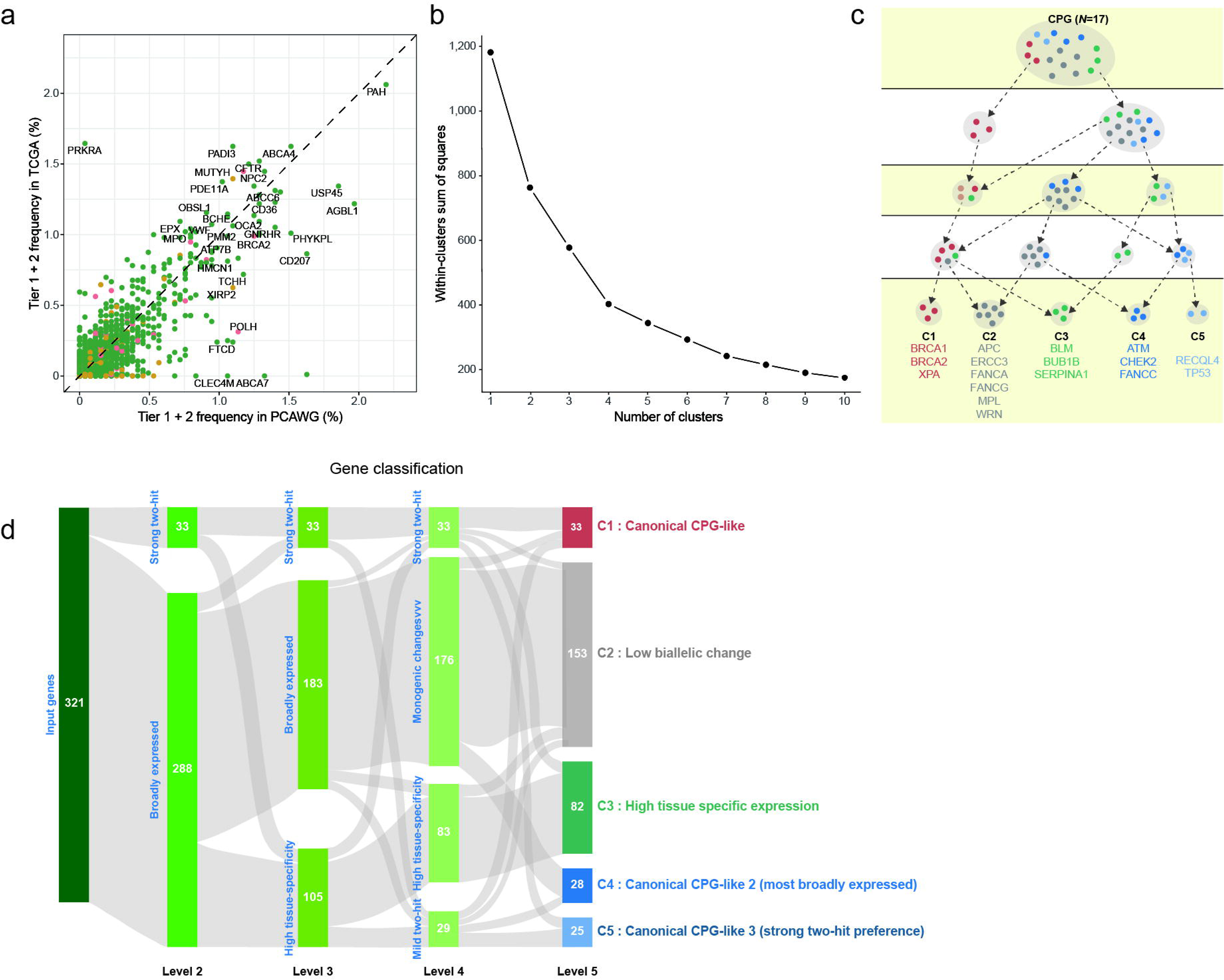
A detailed description of five clusters. (a) Frequency comparison of pathogenic germline variants between PCAWG (x-axis) and TCGA (y-axis) across 3 gene sets (red: CPGs, green: OMIM genes, and yellow: SODs, respectively). (b) A scree plot for determining the optimal number of clusters to use in the trimmed k-means clustering algorithm. (c) Distribution of 17 CPGs across five clusters. (d) Sankey diagram, presenting a classification of CPGs and OMIM genes depending on a cluster.

**Supplementary Figure 5.**
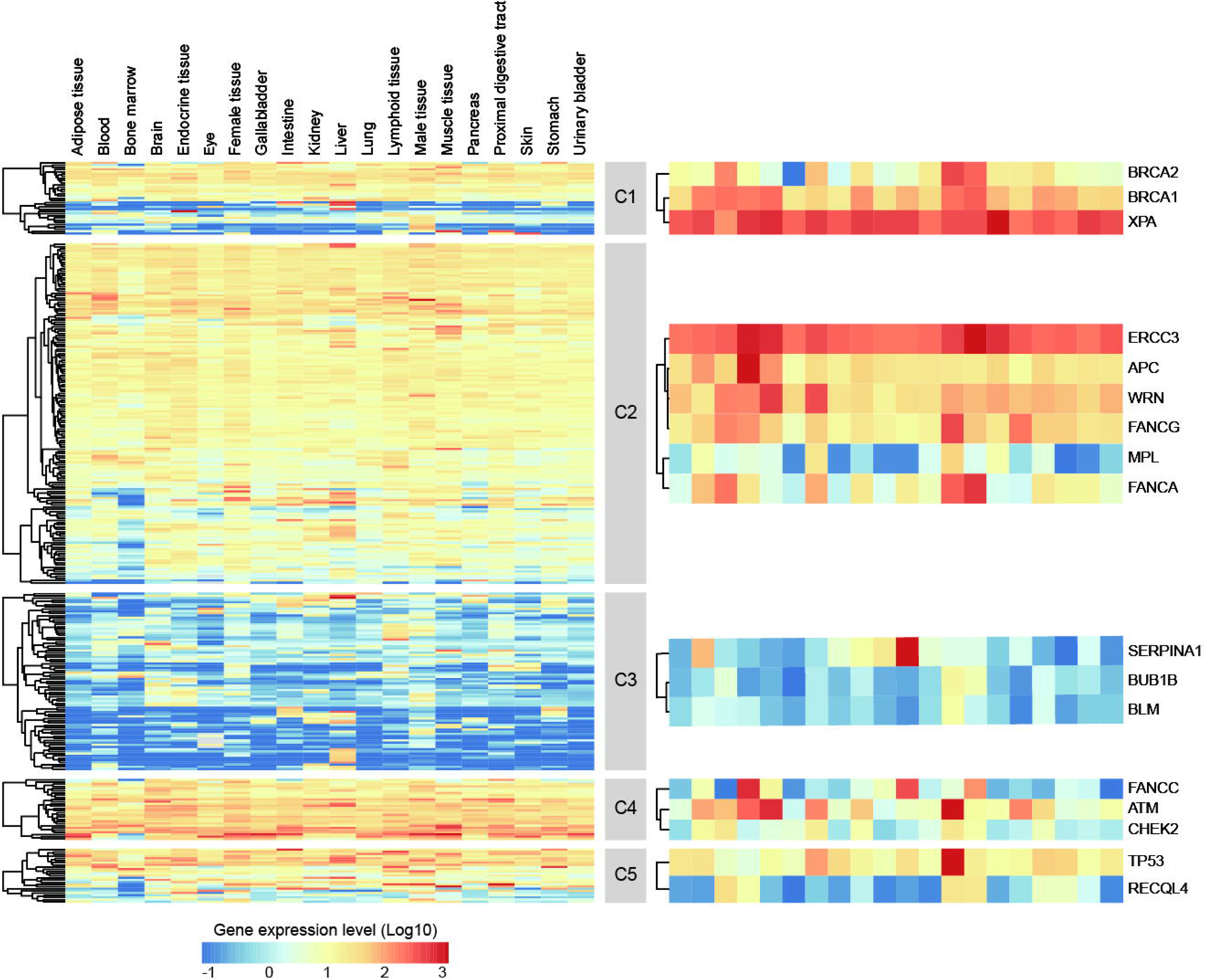
Gene expression level distribution across five clusters. CPGs in each cluster are presented in the right panel. Colors in the heatmap indicate expression level (red: high, blue: low). The columns represent 20 individual tissues from HPA and rows are labelled with individual gene symbols.

**Supplementary Figure 6.**
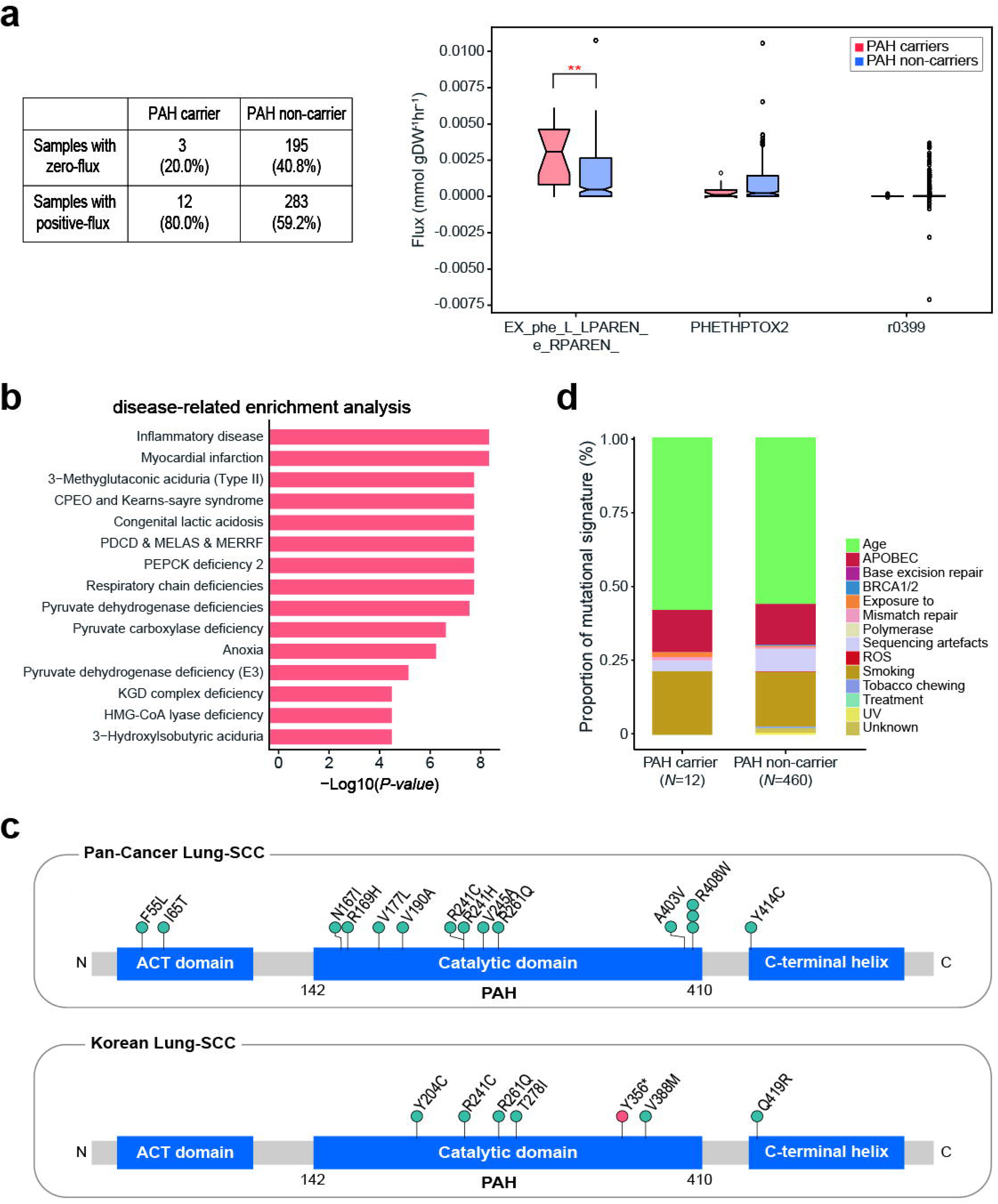
Possible carcinogenic mechanism mediated by PAH. (a) The metabolic simulation results for PAH carriers and non-carriers using genome-scale metabolic models (GEMs) of the corresponding PCAWG-TCGA Lung-SCC samples are summarized in the table (left panel). Box plot presenting the predicted flux of the L-phenylalanine exchange reaction (’EX_phe_L_LPAREN_e_RPAREN_’ in Recon 2M.2) and two additional reactions (’PHETHPTOX2’ and ’r0399’) related to PAH (right panel, ** P < 0.01). The greater the value of the L-phenylalanine exchange reaction means the more active secretion of L-phenylalanine. In addition, the greater values of ’PHETHPTOX2’ and ’r0399’ in PAH non-carriers indicate that L-phenylalanine is more actively converted to L-tyrosine than PAH carriers. (b) Metabolic disease enrichment in PAH carriers compared to non-carrier in Lung-SCC PCAWG-TCGA. Significantly enriched metabolic diseases are presented (P < 0.05, Fisher’s exact test). CPEO: chronic progressive external ophthalmoplegia, PCDC: pyruvate dehydrogenase complex deficiency, MELAS: mitochondrial encephalopathy, lactic acidosis and stroke-like episodes, MERRF: myoclonic epilepsy with ragged-red fibres, PEPCK: Phosphoenolpyruvate carboxykinase, KGD: 2-ketoglutarate dehydrogenase, and HMG-CoA: 3-Hydroxy-3-methylglutaryl-CoA lyase. (c) Lollipop plot showing the rare pathogenic germline variants in PAH using the Lung-SCC (PCAWG-TCGA; top-panel) and Korean Lung-SCC (bottom panel). Blue circles indicate missense variant and red circle represents protein truncating variants. The height of the circle indicates the count of the corresponding variants in the data set (Y-axis). (d) Distribution of somatic mutational signatures in PAH-carriers and PAH non-carriers Lung-SCC PCAWG-TCGA.

## Notes

### Competing Interest Statement

The authors have declared no competing interest.

## References

1. Broca, P.

2. Rahman, N. Realizing the promise of cancer predisposition genes. Nature 505, 302–308, doi:10.1038/nature12981 (2014).

3. Hanahan, D. & Weinberg, R. A. Hallmarks of cancer: the next generation. Cell 144, 646–674, doi:10.1016/j.cell.2011.02.013 (2011).

4. Futreal, P. A. et al. A census of human cancer genes. Nature Reviews Cancer 4, 177–183, doi:10.1038/nrc1299 (2004).

5. Hanahan, D. Hallmarks of Cancer: New Dimensions. Cancer Discov 12, 31–46, doi:10.1158/2159-8290.CD-21-1059 (2022).

6. Chatrath, A., Ratan, A. & Dutta, A. Germline Variants That Affect Tumor Progression. Trends in Genetics 37, 433–443, doi:10.1016/j.tig.2020.10.005 (2021).

7. Hamosh, A., Scott, A. F., Amberger, J. S., Bocchini, C. A. & McKusick, V. A. Online Mendelian Inheritance in Man (OMIM), a knowledgebase of human genes and genetic disorders. Nucleic Acids Res 33, D514–D517, doi:10.1093/nar/gki033 (2005).

8. Milenkovic, I., Blumenreich, S. & Futerman, A. H. GBA mutations, glucosylceramide and Parkinson’s disease. Curr Opin Neurobiol 72, 148–154, doi:10.1016/j.conb.2021.11.004 (2021).

9. Ryland, G. L. et al. Loss of heterozygosity: what is it good for? BMC Med Genomics, 8, 45, doi:10.1186/s12920-015-0123-z (2015).

10. Chan, H. J. et al. SERPINA1 is a direct estrogen receptor target gene and a predictor of survival in breast cancer patients. Oncotarget 6, 25815–25827, doi:10.18632/oncotarget.4441 (2015).

11. Corley, M. et al. An RNA structure-mediated, posttranscriptional model of human α-1-antitrypsin expression. Proc Natl Acad Sci U S A 114, E10244–e10253, doi:10.1073/pnas.1706539114 (2017).

12. Karczewski, K. J. et al. The mutational constraint spectrum quantified from variation in 141,456 humans. Nature 581, 434–443, doi:10.1038/s41586-020-2308-7 (2020).

13. Landrum, M. J. et al. ClinVar: public archive of relationships among sequence variation and human phenotype. Nucleic Acids Res 42, D980–D985, doi:10.1093/nar/gkt1113 (2014).

14. Campbell, P. J. et al. Pan-cancer analysis of whole genomes. Nature 578, 82–93, doi:10.1038/s41586-020-1969-6 (2020).

15. Auton, A. et al. A global reference for human genetic variation. Nature 526, 68–74, doi:10.1038/nature15393 (2015).

16. Huang, K. L. et al. Pathogenic Germline Variants in 10,389 Adult Cancers. Cell 173, 355–370.e314, doi:10.1016/j.cell.2018.03.039 (2018).

17. Gonzalez, H., Hagerling, C. & Werb, Z. Roles of the immune system in cancer: from tumor initiation to metastatic progression. Genes Dev 32, 1267–1284, doi:10.1101/gad.314617.118 (2018).

18. de Visser, K. E., Eichten, A. & Coussens, L. M. Paradoxical roles of the immune system during cancer development. Nature Reviews Cancer 6, 24–37, doi:10.1038/nrc1782 (2006).

19. Decramer, M. & Janssens, W. Chronic obstructive pulmonary disease and comorbidities. Lancet Respir Med 1, 73–83, doi:10.1016/s2213-2600(12)70060-7 (2013).

20. Hatlen, P., Langhammer, A., Forsmo, S., Carlsen, S. M. & Amundsen, T. Bone mass density, fracture history, self-reported osteoporosis as proxy variables for estrogen and the risk of non-small-cell lung cancer--a population based cohort study, the HUNT study: are proxy variables friends or faults? Lung Cancer 81, 39–46, doi:10.1016/j.lungcan.2013.04.001 (2013).

21. Kanehisa, M. & Goto, S. KEGG: kyoto encyclopedia of genes and genomes. Nucleic Acids Res 28, 27–30, doi:10.1093/nar/28.1.27 (2000).

22. Uhlén, M., et al. Proteomics. Tissue-based map of the human proteome. Science 347, 1260419, doi:10.1126/science.1260419 (2015).

23. Blau, N., van Spronsen, F. J. & Levy, H. L. Phenylketonuria. The Lancet 376, 1417–1427, doi:https://doi.org/10.1016/S0140-6736(10)60961-0 (2010).

24. Ryu, J. Y., Kim, H. U. & Lee, S. Y. Framework and resource for more than 11,000 gene-transcript-protein-reaction associations in human metabolism. Proceedings of the National Academy of Sciences 114, E9740–E9749, doi:doi:10.1073/pnas.1713050114 (2017).

25. Song, H. et al. Transcriptional analysis of immune modulatory genes in melanoma treated with PD-1 blockade. bioRxiv, 2020.2012.2020.397000, doi:10.1101/2020.12.20.397000 (2020).

26. Lonsdale, J. et al. The Genotype-Tissue Expression (GTEx) project. Nature Genetics 45, 580–585, doi:10.1038/ng.2653 (2013).

27. Galon, J. & Bruni, D. Approaches to treat immune hot, altered and cold tumours with combination immunotherapies. Nature Reviews Drug Discovery 18, 197–218, doi:10.1038/s41573-018-0007-y (2019).

28. Frederickson, R. M. A New Era of Innovation for CAR T-cell Therapy. Molecular Therapy 23, 1795–1796, doi:https://doi.org/10.1038/mt.2015.205 (2015).

29. Okazaki, T., Iwai, Y. & Honjo, T. New regulatory co-receptors: inducible co-stimulator and PD-1. Curr Opin Immunol 14, 779–782, doi:10.1016/s0952-7915(02)00398-9 (2002).

30. Pavlova, N. N. & Thompson, C. B. The Emerging Hallmarks of Cancer Metabolism. Cell Metab 23, 27–47, doi:10.1016/j.cmet.2015.12.006 (2016).

31. Martinez-Outschoorn, U. E., Peiris-Pagés, M., Pestell, R. G., Sotgia, F. & Lisanti, M. P. Cancer metabolism: a therapeutic perspective. Nature Reviews Clinical Oncology 14, 11–31, doi:10.1038/nrclinonc.2016.60 (2017).

32. Leach, D. R., Krummel, M. F. & Allison, J. P. Enhancement of antitumor immunity by CTLA-4 blockade. Science 271, 1734–1736, doi:10.1126/science.271.5256.1734 (1996).

33. Oscarson, M. Pharmacogenetics of drug metabolising enzymes: importance for personalised medicine. Clin Chem Lab Med 41, 573–580, doi:10.1515/cclm.2003.087 (2003).

34. Burton, B. K. et al. Prevalence of comorbid conditions among adult patients diagnosed with phenylketonuria. Mol Genet Metab 125, 228–234, doi:10.1016/j.ymgme.2018.09.006 (2018).

35. Arbesman, J., Ravichandran, S., Funchain, P. & Thompson, C. L. Melanoma cases demonstrate increased carrier frequency of phenylketonuria/hyperphenylalanemia mutations. Pigment Cell & Melanoma Research 31, 529–533, doi:https://doi.org/10.1111/pcmr.12695 (2018).

36. Li, H. A statistical framework for SNP calling, mutation discovery, association mapping and population genetical parameter estimation from sequencing data. Bioinformatics (Oxford, England) 27, 2987–2993, doi:10.1093/bioinformatics/btr509 (2011).

37. Scheinin, I. et al. DNA copy number analysis of fresh and formalin-fixed specimens by shallow whole-genome sequencing with identification and exclusion of problematic regions in the genome assembly. Genome Research 24, 2022–2032, doi:10.1101/gr.175141.114 (2014).

38. Amemiya, H. M., Kundaje, A. & Boyle, A. P. The ENCODE Blacklist: Identification of Problematic Regions of the Genome. Scientific Reports 9, 9354, doi:10.1038/s41598-019-45839-z (2019).

39. Derrien, T. et al. Fast Computation and Applications of Genome Mappability. PLOS ONE 7, e30377, doi:10.1371/journal.pone.0030377 (2012).

40. Wang, K., Li, M. & Hakonarson, H. ANNOVAR: functional annotation of genetic variants from high-throughput sequencing data. Nucleic Acids Res 38, e164–e164, doi:10.1093/nar/gkq603 (2010).

41. McLaren, W. et al. The Ensembl Variant Effect Predictor. Genome Biology 17, 122, doi:10.1186/s13059-016-0974-4 (2016).

42. Purcell, S. et al. PLINK: a tool set for whole-genome association and population-based linkage analyses. Am J Hum Genet 81, 559–575, doi:10.1086/519795 (2007).

43. Zhang, J. et al. Germline Mutations in Predisposition Genes in Pediatric Cancer. New England Journal of Medicine 373, 2336–2346, doi:10.1056/NEJMoa1508054 (2015).

44. Martínez-Jiménez, F. et al. A compendium of mutational cancer driver genes. Nature Reviews Cancer 20, 555–572, doi:10.1038/s41568-020-0290-x (2020).

45. Lawrence, M. S. et al. Discovery and saturation analysis of cancer genes across 21 tumour types. Nature 505, 495–501, doi:10.1038/nature12912 (2014).

46. Dietlein, F. et al. Identification of cancer driver genes based on nucleotide context. Nature Genetics 52, 208–218, doi:10.1038/s41588-019-0572-y (2020).

47. Durinck, S., Spellman, P. T., Birney, E. & Huber, W. Mapping identifiers for the integration of genomic datasets with the R/Bioconductor package biomaRt. Nature Protocols 4, 1184–1191, doi:10.1038/nprot.2009.97 (2009).

48. Liberzon, A. et al. Molecular signatures database (MSigDB) 3.0. Bioinformatics 27, 1739–1740, doi:10.1093/bioinformatics/btr260 (2011).

49. Dong, X., Hao, Y., Wang, X. & Tian, W. LEGO: a novel method for gene set over- representation analysis by incorporating network-based gene weights. Scientific Reports 6, 18871, doi:10.1038/srep18871 (2016).

50. Li, Y. et al. Patterns of somatic structural variation in human cancer genomes. Nature 578, 112–121, doi:10.1038/s41586-019-1913-9 (2020).

51. Goldman, M. J. et al. A user guide for the online exploration and visualization of PCAWG data. Nature Communications 11, 3400, doi:10.1038/s41467-020-16785-6 (2020).

52. Dentro, S. C. et al. Characterizing genetic intra-tumor heterogeneity across 2,658 human cancer genomes. Cell 184, 2239–2254.e2239, doi:https://doi.org/10.1016/j.cell.2021.03.009 (2021).

53. Fritz, H., García-Escudero, L. A. & Mayo-Iscar, A. tclust: An R Package for a Trimming Approach to Cluster Analysis. Journal of Statistical Software 47, 1–26, doi:10.18637/jss.v047.i12 (2012).

54. Grossman, R. L. et al. Toward a Shared Vision for Cancer Genomic Data. New England Journal of Medicine 375, 1109–1112, doi:10.1056/NEJMp1607591 (2016).

55. Pang, Z. et al. MetaboAnalyst 5.0: narrowing the gap between raw spectra and functional insights. Nucleic Acids Res 49, W388–W396, doi:10.1093/nar/gkab382 (2021).

56. Gu, C., Kim, G. B., Kim, W. J., Kim, H. U. & Lee, S. Y. Current status and applications of genome-scale metabolic models. Genome Biology 20, 121, doi:10.1186/s13059-019-1730-3 (2019).

57. Dobin, A. et al. STAR: ultrafast universal RNA-seq aligner. Bioinformatics 29, 15–21, doi:10.1093/bioinformatics/bts635 (2012).

58. Li, B. & Dewey, C. N. RSEM: accurate transcript quantification from RNA-Seq data with or without a reference genome. BMC Bioinformatics 12, 323, doi:10.1186/1471-2105-12-323 (2011).

59. Subramanian, A. et al. Gene set enrichment analysis: A knowledge-based approach for interpreting genome-wide expression profiles. Proceedings of the National Academy of Sciences 102, 15545–15550, doi:10.1073/pnas.0506580102 (2005).

60. Alexandrov, L. B. et al. The repertoire of mutational signatures in human cancer. Nature 578, 94–101, doi:10.1038/s41586-020-1943-3 (2020).

61. Poplin, R. et al. Scaling accurate genetic variant discovery to tens of thousands of samples. bioRxiv, 201178, doi:10.1101/201178 (2018).

